# Whole forest in a pouch? Methods converge in uncovering wood ants’ fungal and bacterial microbiota

**DOI:** 10.64898/2026.03.24.713951

**Authors:** Igor Siedlecki, Michał Kochanowski, Izabela Bąk, Michał Kolasa, Mateusz Buczek, Karol H. Nowak, Zuzanna Błocka, Zuzanna Płoszka, Julia Pawłowska, Piotr Łukasik, Marta Wrzosek

## Abstract

Despite their importance for individual fitness and population processes, the microbiota of many ecologically significant insects remains poorly explored. Even less is known about the interactions between microbial communities inhabiting insects and their surrounding environment. Ant infrabuccal pockets (IBPs), representing the interface between the digestive tract and the external environment, provide an opportunity to study these interactions. Here, we aimed to characterize ant–microbial interaction networks in the forest floor by profiling fungal and bacterial communities associated with the IBP of *Formica polyctena* ants, known as ecosystem engineers in temperate forests. We used direct microscopy, culturing, and sequencing amplicons of ITS1, ITS2, 18S rRNA marker regions to describe fungal communities, and 16S rRNA metabarcoding to characterize bacterial communities. Classical methods combined with a multi-marker amplicon sequencing allowed for a comprehensive description of the IBP microbiota. Fungal communities consistently contained representatives of 15 ecologically diverse genera, including insect-associated yeasts and primarily saprotrophic or endophytic fungi. Bacterial communities were dominated by genera previously reported from ant guts, mainly *Bacilli* and *Alphaproteobacteria*, and showed greater stability among ant colonies than fungal communities. Further studies on red wood ants’ IBP microbiota would enhance our understanding of their role in shaping ecological networks in forest ecosystems.

## 1. Introduction

Interspecies interactions play a critical role in determining biodiversity and the stability of ecosystems, as they strongly affect organisms’ ecological niches (Duffy et al. 2007; López-Pujol, 2011; Wisz et al. 2013). In such interactions, microorganisms commonly act as symbionts residing in larger hosts (Gibson and Hunter, 2010). However, while bacteria and microscopic fungi have long been known for their roles in key ecosystem processes (Barton and Northup, 2011; Dix, 2012), their critical roles in many aspects of higher organisms’ biology, including nutrition, reproduction, and defense, have been recognized only recently (Bahrndorff et al. 2016; Jordan and Tomberlin, 2021; Łukasik and Kolasa, 2024; McFall-Ngai et al. 2013; Vandenkoornhuyse et al. 2015).

Although ants (*Hymenoptera*: *Formicidae*) are among the most abundant and broadly distributed groups of insects, playing a significant role in ecosystem functioning (Del Toro et al. 2012; Folgarait, 1998; Hölldobler and Wilson, 1990), our understanding of their interactions with fungal and bacterial communities remains incomplete (Gibson and Hunter, 2010; Moreau, 2020). Aside from fungus-farming ants that maintain mutualistic relationships with their symbiotic fungal cultivars (Schultz et al. 2024), and ants inhabiting domatia or forming cardboard nests in symbiosis with the black yeasts (*Ascomycota*: *Chaetothyriales*) (Blatrix et al. 2012; Nepel et al. 2014; Voglmayr et al. 2011), systematic efforts to characterize non-pathogenic fungal associations within this important insect family are limited. Similarly, only a few ant clades can be described as having relatively well-understood bacterial communities. For example, carpenter ants are known to host mutualistic intracellular bacteria in the genus *Blochmannia* (Boursaux-Eude and Gross, 2000; Feldhaar et al. 2007), turtle ants harbor diverse but conserved gut bacterial communities (Hu et al. 2018), and army ants associate with specialized gut-occupying *Entomoplasmatales* and *Firmicutes* (Funaro et al. 2011; Łukasik et al. 2017). On the other hand, some ant taxa may lack abundant specialized microbiota (Hammer et al. 2019; Sanders et al. 2017). However, until now, most studies on ant-associated microbiota have either relied on fully homogenized individuals or focused on dissected gasters and guts, neglecting other parts of the digestive tract that may harbor distinct microbial communities (Callegari et al. 2021; Lanan et al. 2016).

The ants’ infrabuccal pocket (IBP) is a part of their digestive tract that has the potential to provide particularly broad insights into ant biological interactions. This pouch-like organ, located in the head, serves as a reservoir for filtered particles collected during feeding and grooming. Inside the IBP, these particles are formed into a pellet, which is periodically ejected (Eisner and Happ, 1962; Richter and Economo, 2023). In addition to filtering particles, the infrabuccal pocket is hypothesized to play a role in food digestion (Hansen et al. 1999; Richter and Economo, 2023) and disease protection (Little et al. 2005). Despite numerous studies focused on the morphology of this organ (e.g., Richter et al. 2020; Wang et al. 2019; Wang et al. 2022), very little is known about the composition, function, and potential origins of the microbial communities present in the IBP.

To date, studies on the IBP’s bacterial microbiome have revealed a multitude of bacteria present in this organ in *Camponotus modoc*, *Paraponera clavata*, and *Cephalotes* ants (Hansen et al. 1999; Moreau and Rubin, 2017; Rodrigues, 2016). Additionally, differences in bacterial communities between the IBP and other parts of the ants’ digestive tract have been reported for three species of *Formica*, *Lasius niger*, and *P. clavata* (Moreau and Rubin, 2017; Zheng et al. 2022). The composition of the IBP bacterial community is suggested to be linked to diet, as lactic acid bacteria were abundantly observed in ants that primarily feed on aphid honeydew (Zheng et al. 2022). Compared with bacteria, far fewer studies have focused on fungal communities of the IBP, despite the early findings by Wheeler and Bailey (1920) of fungal material inside the pellets of numerous ant species. To date, mutualistic fungi have been observed to be transported in the IBP of queens establishing new colonies only in leaf-cutting and domatia-occupying ants (Hölldobler and Wilson, 1990; Mayer et al. 2018). Additionally, studies on *Camponotus* ants showed that saprotrophs typical of ant nests and surrounding soil were commonly isolated from the IBP (Clark, 2002; Mankowski and Morrell, 2004). However, apart from these few examples, information on the diversity and nature of fungal communities associated with the IBP in other ants’ taxa remains unknown.

Red wood ants (RWA, *Formica rufa* group) are among the most ecologically and economically significant ant clades, with pronounced infrabuccal pockets (Richter et al. 2020; Stockan and Robinson, 2016). RWA are considered key players in forest ecosystems, and are commonly found in Eurasian temperate forests, where they construct large prominent mounds composed of collected plant material (Czechowski, 2012; Frouz and Jilková, 2008; Risch et al. 2005; Stockan and Robinson, 2016). They interact with a wide range of animals and plants, including engaging in mutualistic interactions with sap-sucking insects (Stockan and Robinson, 2016), myrmecochory (Gorb and Gorb, 1999), hosting myrmecophiles (Parmentier et al. 2014), preying on folivores (Karhu and Neuvonen, 1998), and competing with ground beetles (Hawes et al. 2013). However, knowledge of RWA interactions with microscopic fungi and bacteria is still limited. Recent studies using DNA metabarcoding characterized bacterial communities in fully homogenized RWA individuals, showing they are mostly composed of *Anaplasmataceae*, *Acetobacteraceae*, and *Lactobacillaceae* (Jackson et al. 2023; Kaczmarczyk-Ziemba et al. 2020; Sinotte et al. 2024). Regarding fungal communities associated with RWA, culture-dependent methods revealed a significantly different yeast community in *Formica aquilonia* ants and their mounds compared with the surrounding soil, with particularly abundant representatives of *Debaryomycetaceae* (Maksimova et al. 2016). Additionally, distinct communities of *Mucoromycota* in the mounds of *F. polyctena* have been reported (Siedlecki et al. 2024), and *Penicillium* species were abundantly isolated from the cadavers of these ants (Siedlecki et al. 2021). However, there is little molecular data on these fungal associations, and no studies have simultaneously addressed bacterial and fungal communities associated with RWA.

The goal of the study was to characterize fungal and bacterial communities associated with the infrabuccal pockets of RWA, specifically in *Formica polyctena*. To obtain the most complete overview of fungal communities in ant infrabuccal pellets, we used three parallel but fundamentally different methods (direct microscopy, culturing, and multi-marker DNA metabarcoding). To characterize the bacterial community, we used 16S rRNA metabarcoding. We aimed to identify the predominant taxa present, compare fungal communities obtained using different methods, and assess the stability and ecological specificity of the IBP microbial communities.

## 2. Methods

### 2.0. Experimental overview

We first collected *Formica polyctena* ants and isolated pellets from their infrabuccal pockets (IBP) (Fig. 1A). To assess the fungal community composition of the IBP, we used three parallel methods: (a) direct microscopy, (b) culturing supplemented with barcoding of representative strains (ITS and LSU regions), and (c) multi-marker DNA metabarcoding (ITS1, ITS2, and 18S rRNA regions) (Fig. 1B). To characterize the bacterial community of the IBP, we used 16S rRNA metabarcoding (Fig. 1C). Finally, we assessed the stability of the microbiota among colonies and its ecological specificity by examining the environmental distribution of fungal and bacterial amplicon sequence variants (ASVs) abundantly detected in the pellets (core ASVs) through comparisons with sequences stored in the NCBI database (Fig. 1D).

**Figure 1.**
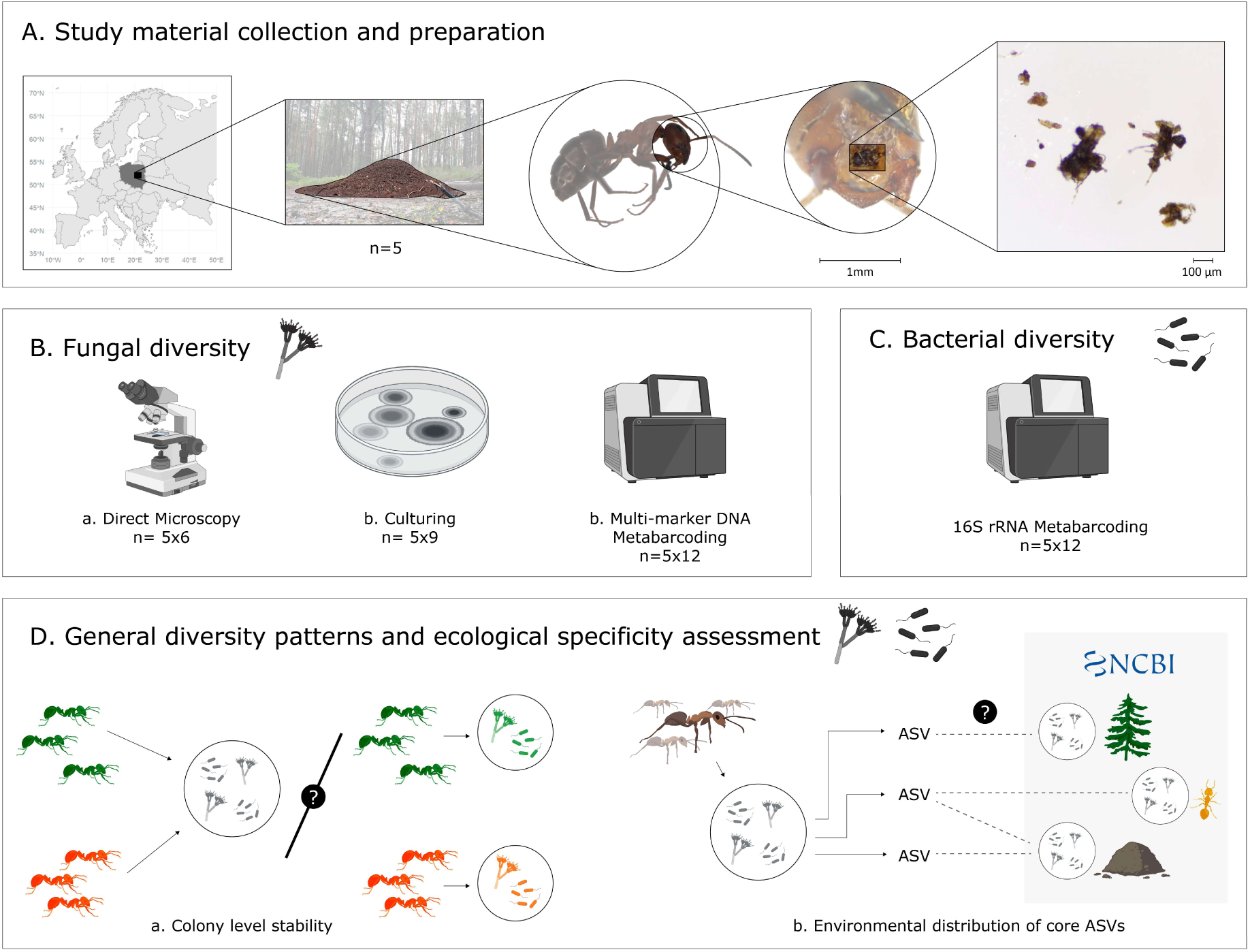
Experimental overview. **A**. The origin of samples; **B-C**. Approaches for the study of fungal and bacterial diversity, and sample sizes. **D**. Conceptual approaches to the interpretation of patterns. The photos in the upper part show *Formica polyctena* ants’ mound, worker, and head with the exposed infrabuccal pellet, which is further shown in detail.

### 2.1. Study material collection and preparation

To acquire the contents of ants’ IBPs, we collected *F. polyctena* workers from five colonies (CHO, KOR, FAL, ZAB, and ZIE) located in distinct pine forests in central Poland, separated by a minimum distance of 10 km, between 15 and 27 August 2019 (Supplementary Table ST.1). We identified the species using the key by Czechowski (2012). We collected at least 30 ants per colony, along with nest material, from within the mounds. In the laboratory, we separated the ants from the nest material. Workers designated for fungal culturing were euthanized by freezing at −22°C for 25 minutes and dissected on the same day. The remaining ants, used for direct microscopy and metabarcoding, were stored at −80°C until further processing. We isolated the contents of the infrabuccal pockets under sterile conditions, following the procedures described by Clark (2002) and shown in Appendix A.1. Ants were collected under permission from the Regional Directorate for Environmental Protection (RDOŚ) in Warsaw (permit number: WPN-I.6401.180.2019.AZ.19).

### 2.2. Direct Microscopy

We placed infrabuccal pellets from 30 ants (6 workers per colony) separately in a drop of lactophenol blue on a glass slide. Using a coverslip, we gently spread the material and inspected it at 100× - 600× magnification under a Nikon ECLIPSE Ni-U light microscope. Imaging was performed with a Nikon DS-Ri2 camera and NIS Elements D 5.11.00 software. We grouped the observed organisms or organismal parts into six general taxonomic categories: bacteria, fungal matter, animal tissue, plant tissue, lichens/algae, and slime molds (see Appendix A.2 for example photographs). Additionally, we assessed the abundance of fungal and bacterial material using the classification system of Wheeler and Bailey (1920): Few – very small amounts of material, either absent or only sporadically visible; Numerous – moderate amounts of material, visible in part of the microscopic fields of view, covering no more than half of the field; Very Numerous – large amounts of material, visible in most or all microscopic fields, occupying a significant portion of the view.

We further classified fungal structures into morphological types using taxonomic keys and monographs (e.g., Breitenbach and Kränzlin, 1984; Læssøe and Petersen, 2019; Seifert et al. 2011; Skirgiełło et al. 1979). We distinguished morphotypes primarily based on differences in shape, size, clustering, number of cells, color, and wall ornamentation of reproductive structures. In some cases, we distinguished morphotypes based on specific hyphal structures (e.g., *Entomortierella*) or presence of characteristic fungal structures within other organisms (e.g., *Rozellomycota* in pine pollen). We measured fungal structures using the Fiji image processing package in ImageJ 2.15.1 (Schindelin et al. 2012). We named morphotypes, as collective categories, under the best-fitting taxonomic name with the suffix ‘-like’. To determine the prevalence of different morphotypes, we recorded the presence of each morphotype in every studied pellet. Distinguished morphotypes, along with representative photographs, are available in Appendix A.3. Direct microscopy datasets used for further analysis and visualization are available in Supplementary Tables ST.2-3.

### 2.3. Culture-dependent diversity

#### 2.3.1. Isolation of Fungi

For fungal culturing, we isolated 45 infrabuccal pellets from nine workers per colony across five colonies. Each pellet was placed into a centrifuge tube containing 0.6 mL of 0.9% saline buffer and vortexed until it disintegrated. We evenly distributed 0.2 mL of the resulting suspension onto three Petri dishes containing Sabouraud Dextrose 4% Agar (SDA) supplemented with 0.05 g/L chloramphenicol. Plates were incubated at 20°C in darkness and monitored daily. Slow-growing strains at risk of being overgrown by fast-growing strains were transferred to fresh sterile SDA medium. After 7 days of incubation, we moved the Petri dishes to 4°C and stored them for up to 14 days, until further processing.

#### 2.3.2. Identification of isolates

We identified the isolates based on molecular and morphological characteristics, as described previously (Siedlecki et al. 2024). Briefly, we observed macroscopic features using a stereoscopic microscope and examined the strains’ micromorphology under a light microscope. For sporulating fungi, we grouped colonies with similar morphology into distinct morphotypes using monographs and keys (Seifert et al. 2011; Skirgiełło et al. 1979). We preserved two representative strains from each morphotype as axenic cultures. Additionally, we treated non-sporulating strains of *Basidiomycota* and yeast colonies as separate general categories and counted them without further identification.

We extracted total genomic DNA from all representative strains. Molecular identification was based primarily on the internal transcribed spacer region (ITS), supplemented with the large subunit nuclear ribosomal DNA (LSU) marker for strains for which ITS amplification failed. Primer sequences and amplification conditions are listed in Appendix A.4. Purified PCR products were sequenced using the Sanger method by Genomed S.A. (Warsaw, Poland) and NexBio (Lublin, Poland). We assembled forward and reverse reads into contigs and manually verified them using the DNA Subway software (Williams et al. 2014). We compared the consensus sequences using the BLASTn algorithm (Altschul et al. 1990) against the core_nt database at NCBI, restricted to sequences of type and reference strains deposited in recognized public culture collections (ncbi.nlm.nih.gov, accessed on 13 November 2024).

We assigned species-level names to morphotypes when morphology-based identification aligned with top BLAST hits, meeting the following criteria: (1) sequence coverage >90%, (2) sequence identity >97% for ITS and >98% for LSU, and (3) sequence divergence from the closest related species >0.7% for ITS and >0.3% for LSU, with both representative strains assigned to the same species. Otherwise, a higher-level taxonomic rank was used. Additionally, we assigned UNITE Species Hypotheses (SH; 1.5% threshold) number to sequences using SH matching tool on the PlutoF platform (https://plutof.ut.ee/en, accessed on 13 November 2024), with Run ITSx and substring dereplication enabled (Abarenkov et al. 2022). A list of representative strains with identification details, voucher numbers, and generated sequences is provided in Supplementary Table ST.4.

### 2.4. Multi-marker DNA metabarcoding

#### 2.4.1. DNA extraction, library preparation and sequencing

We processed the samples using modified protocols detailed by Buczek et al. (2024). For multi-marker DNA metabarcoding, we used 60 infrabuccal pellets isolated from 12 individual workers from each of the five colonies. Each pellet was suspended in 30 µL of PCR-grade water and placed in 2 mL tubes containing 200 µL of lysis buffer (0.4 M NaCl, 10 mM Tris-HCl, 2 mM EDTA, pH 8, 2% SDS) (Aljanabi and Martinez, 1997; Vesterinen et al. 2016), including 5 µL of lyticase (A&A Biotechnology, Poland). After adding 2.8 mm and 0.5 mm ceramic beads, samples were homogenized using an Omni Bead Raptor Elite homogenizer for two 30-second cycles at a speed of 5 m/s, followed by a two-hour incubation at room temperature. Subsequently, 5 µL of proteinase K (Thermo Fisher Scientific Inc., USA) was added to each sample, and samples were incubated for an additional two hours at 55°C. Forty µL of homogenate from each tube was purified using 80 µL of SPRI beads and eluted in 20 µL of TE buffer.

For each sample, we prepared two amplicon libraries targeting different marker regions. The first library targeted four combined regions: the V4 region of the 16S rRNA bacterial gene, a portion of the insect mitochondrial cytochrome oxidase I (COI) gene, and two fungal markers: the ITS1a and ITS2 regions of the nuclear ribosomal internal transcribed spacer. The second library targeted a c.a. 350-bp fragment of the fungal 18S rRNA gene DNA (hereafter referred to as 18S). We followed a two-step PCR protocol. In the first step, marker regions were amplified using a mixture of template-specific primer pairs with Illumina adapter tails. Products were verified on a 2.5% agarose gel, cleaned with SPRI beads, and used as templates for the second indexing PCR, in which Illumina adapters were completed and unique indexes were added to each sample. Primer sequences and reaction conditions are listed in Appendix A.4. During all steps, we included positive and negative controls to verify accuracy. After verifying the libraries on an agarose gel, we pooled them based on band intensity and sequenced alongside other amplicon libraries on P1 and P2 flow cells (600-cycle) using an Illumina NextSeq 2000 at the Malopolska Centre of Biotechnology, Jagiellonian University.

#### 2.4.2. Bioinformatic analysis

We performed the bioinformatic analysis on the Jagiellonian University’s Unix cluster using a pipeline combining custom Python scripts with established bioinformatics tools available at https://github.com/IgorSiedlecki/Microbial-diversity-of-Formica-polyctena-infrabuccal-pockets. First, we split the amplicon data into bins corresponding to the targeted regions based on the primer sequences. For each bin separately, we assembled quality-filtered forward and reverse reads into contigs using PEAR with a minimum Phred score of 30 (Zhang et al. 2014). Next, we de-replicated the contigs (Rognes et al. 2016) and denoised them using UNOISE3, with the - minsize option set to 8 (Edgar, 2016). We processed each library separately to avoid losing information about genotypes present only in one or a few individuals, which could occur during denoising of the entire dataset at once (Prodan et al. 2020). We screened the sequences for chimeras using USEARCH (Edgar, 2010) and assigned taxonomy using the SINTAX algorithm (Edgar, 2010) and customized reference databases (see Appendix A.5). We accepted phylum-level assignments with a probability ≥0.75, and assignments at other taxonomic levels with a probability ≥0.9. We assigned taxonomic names to ASVs up to the genus level but omitted species-level assignments. Additionally, since the SILVA and UNITE taxonomy reference databases are based on differing taxonomic backbones, we adjusted taxonomy assignments for 18S rRNA reads to enable comparison of fungal diversity using the ITS marker gene (see Appendix A.6). To complement taxonomy assignment for ASVs, we compared genotypes with a maximum relative abundance of >1% in any library and not classified to the genus level with data available in NCBI GenBank (ncbi.nlm.nih.gov, accessed on 13 November 2024) using the BLASTn algorithm (Altschul et al. 1990) (see Appendix A.7). Finally, we clustered the sequences at a 97% identity level using the UPARSE-OTU algorithm implemented in USEARCH. We produced tables with two levels of classification: ASVs, describing genotypic diversity, and OTUs (Operational Taxonomic Units) clustering ASVs based on a similarity threshold.

In order to clean the datasets from laboratory and reagent-derived contaminants (Salter et al. 2014), we implemented a custom filter (Łukasik et al. 2017) based on relative abundances of ASVs for fungi and OTUs for bacteria in experimental samples and negative controls (see Appendix A.8). For the fungal 18S dataset, we applied an additional filtering step informed by the ITS marker gene results. Because *Cladosporium* genotypes occurred abundantly in both experimental samples and blanks, and ITS data indicated that only a subset represented reagent contamination, we removed only a corresponding fraction of *Cladosporium* 18S reads (see Appendix A.9). Next, we used taxonomy classification information to remove non-bacterial or non-fungal reads, depending on the marker region. From 16S rRNA data, we excluded ASVs classified as chloroplasts, mitochondria, *Archaea*, or chimeras. From ITS1 and ITS2 data, we retained only genotypes assigned to Fungi with a probability ≥0.75. For 18S rRNA, we retained only genotypes assigned to fungal phyla with a probability ≥0.5. Finally, per each marker gene we removed libraries with fewer than 1000 reads. Because only 8% of libraries (5 IBPs) in the COI dataset met this threshold, and most reads corresponded to ants, COI data was excluded from the analysis in this paper. The decontaminated datasets with adjusted taxonomy, used for statistical analysis and visualizations, are available in Supplementary Tables ST.7–10.

### 2.5. Comparison of methods to study fungal diversity

To compare independent methods for studying fungal diversity, we compared the number of fungal genera obtained from the IBPs using different approaches. To verify the consistency of the main results across methods, we assigned the most prevalent taxa identified by culturing and multi-marker metabarcoding to the seven most prevalent fungal morphotypes observed by direct microscopy. To compare molecular diversity between culturing and metabarcoding approaches, we calculated the percentage of matching sequences (>99% identity and >75% overlap) between ITS Sanger sequences obtained from cultured strains, and ITS1 and ITS2 ASVs obtained from multi-marker metabarcoding.

### 2.6. Determination of general diversity patterns and ecological specificity of described communities

For the diversity and specificity analyses, we used two metabarcoding datasets: 16S rRNA for bacteria, and for fungi ITS2 - deemed the most informative. To determine the general diversity patterns in the studied communities, we calculated ASV richness using the *specnumber* function from the ‘vegan’ package (Oksanen et al. 2025). We tested for differences in ASV richness among colonies using the Kruskal–Wallis test, implemented in R with the *kruskal.test* function. Pairwise comparisons were conducted using Dunn’s test with Holm correction via the *dunnTest* function from the ‘FSA’ package (Ogle et al. 2015). We analyzed differences in fungal community composition between sampling sites using permutational multivariate analysis of variance (PERMANOVA) with the *adonis2* function (Bray–Curtis distance, 999 permutations) from the ‘vegan’ package (Oksanen et al. 2025). Pairwise comparisons of microbial community differences were performed using the *pairwise.adonis* function from the ‘pairwiseAdonis’ library (Arbizu, 2020). We assessed the homogeneity of variances within and between groups using the *betadisper* function from the ‘vegan’ package. All analyses were conducted in R version 4.4.1 and RStudio version 2024.09.0.

To determine the ecological specificity of communities, we compared sequences of the 20 most abundant ITS2 and 16S rRNA ASVs (hereafter “core ASVs”) against the NCBI nucleotide database (accessed on 27 January 2026) using BLASTn (Altschul et al. 1990). For each core ASV, we determined its environmental distribution based on the isolation source information available in the descriptions of the first 10 sequences with 100% identity. We distinguished 13 environmental distribution categories, listed in Supplementary Table ST.12. If a genotype matched sequences isolated from various types of sources, we assigned multiple environmental distribution categories.

### 2.7. Data Visualization

We created all heatmaps using Processing 3.5.4 (Reas and Fry, 2006). We performed other data visualizations using R version 4.4.1 and RStudio version 2024.09.0. For fungi, we generated a Venn diagram illustrating shared and unique genera noted for different gene markers in the multi-marker metabarcoding approach using the *venn.diagram* function from the ‘VennDiagram’ package (Chen, 2022). We created all other figures using the ‘ggplot2’ package (Wickham et al. 2024). Additionally, we used non-metric multidimensional scaling (NMDS) ordinations based on Bray–Curtis distances (calculated using the *metaMDS* function) to visualize bacterial (16S rRNA) and fungal (ITS2) community composition. Before analysis, the data were square-root transformed and subjected to Wisconsin double standardization.

## 3. Results

### 3.1. The biological and fungal diversity assessed using microscopy

Within 30 *F. polyctena* IBPs, we observed a variety of organisms (Fig. 2A; Appendix A.2). All pellets contained bacteria and fungi, both typically scored as ‘numerous’ or ‘very numerous’, and fragments of plant tissue or pollen grains. Approximately half of the pellets contained remains of animals or algae. Additionally, we observed slime mold spores in about one-quarter of the pellets (Fig. 2A). We distinguished a total of 101 fungal morphotypes (Appendix A.3), with an average of 20 morphotypes per pellet (range: 3–37). Colonies differed in the number of morphotypes detected (Kruskal-Wallis test: χ² = 19.264, df = 4, p = 0.0007), with significantly fewer morphotypes observed in IBPs from colony CHO than in KOR, FAL, and ZAB (Wilcoxon test with continuity correction, Holm adjustment, p < 0.05). Twelve morphotypes were observed in at least half of all IBPs (Fig. 2B). The most prevalent morphotypes were *Cladosporium/Cladophialophora*-like, bacteria-like microspores, and dark-hyphae morphotypes, with the first two occurring in all pellets and the last one in 93% of the pellets.

**Figure 2.**
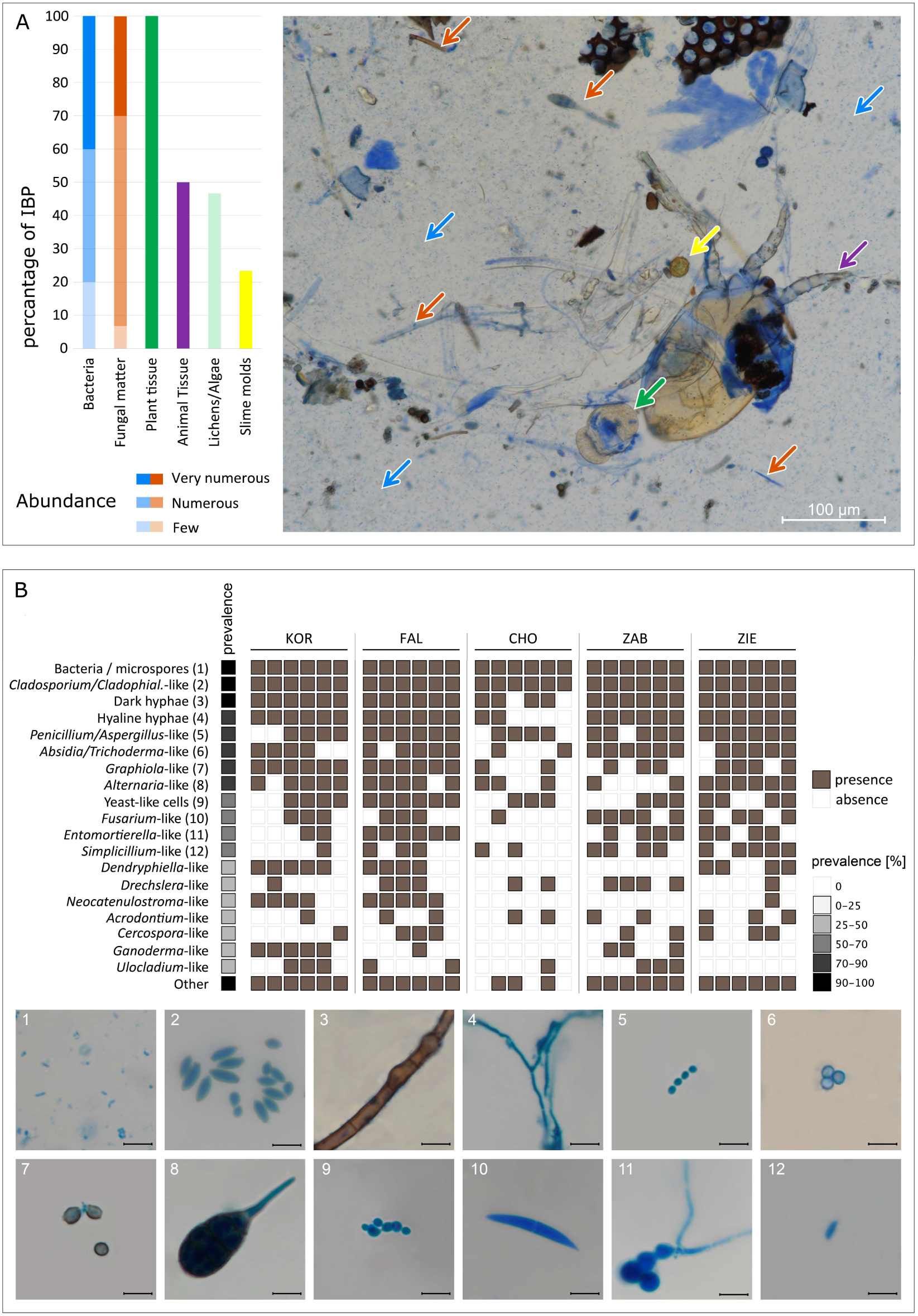
Biological diversity within ants’ infrabuccal pellets, assessed by direct microscopy. **A.** The prevalence and estimated abundance of broad taxonomic categories, and their representatives in an example pellet - indicated using an arrow of the same color as the taxonomic label. **B**. The prevalence of the most common fungal morphotypes in IBPs from across processed specimens representing five colonies, with morphotypes appearing in <30% IBPs grouped together as “Others”. Prevalence is calculated as the percentage of IBPs in which a given taxon was recorded. Photograph numbers correspond to morphotype numbers in the upper portion of panel B. Scale bar: 10 μm.

### 3.2. Diversity of cultured fungi

We isolated a total of 5945 microbial colonies from 45 IBPs. Despite the use of bacteriostatic chloramphenicol, this number included 3930 bacterial colonies, which grew from 69% of the pellets. Fungal colonies grew from all pellets, totaling 2015 colony-forming units (CFUs), with an average of 45 fungal CFUs per pellet (range: 1–668, SD: 110). We observed no significant differences in the number of isolated fungal CFUs among ant colonies (Fig. 3A). For 22% of the pellets, we observed limited fungal growth (<5 CFUs) (Fig. 3A). We also noted that 6% of fungal colonies became completely overgrown by other strains, leaving them unidentified.

**Figure 3.**
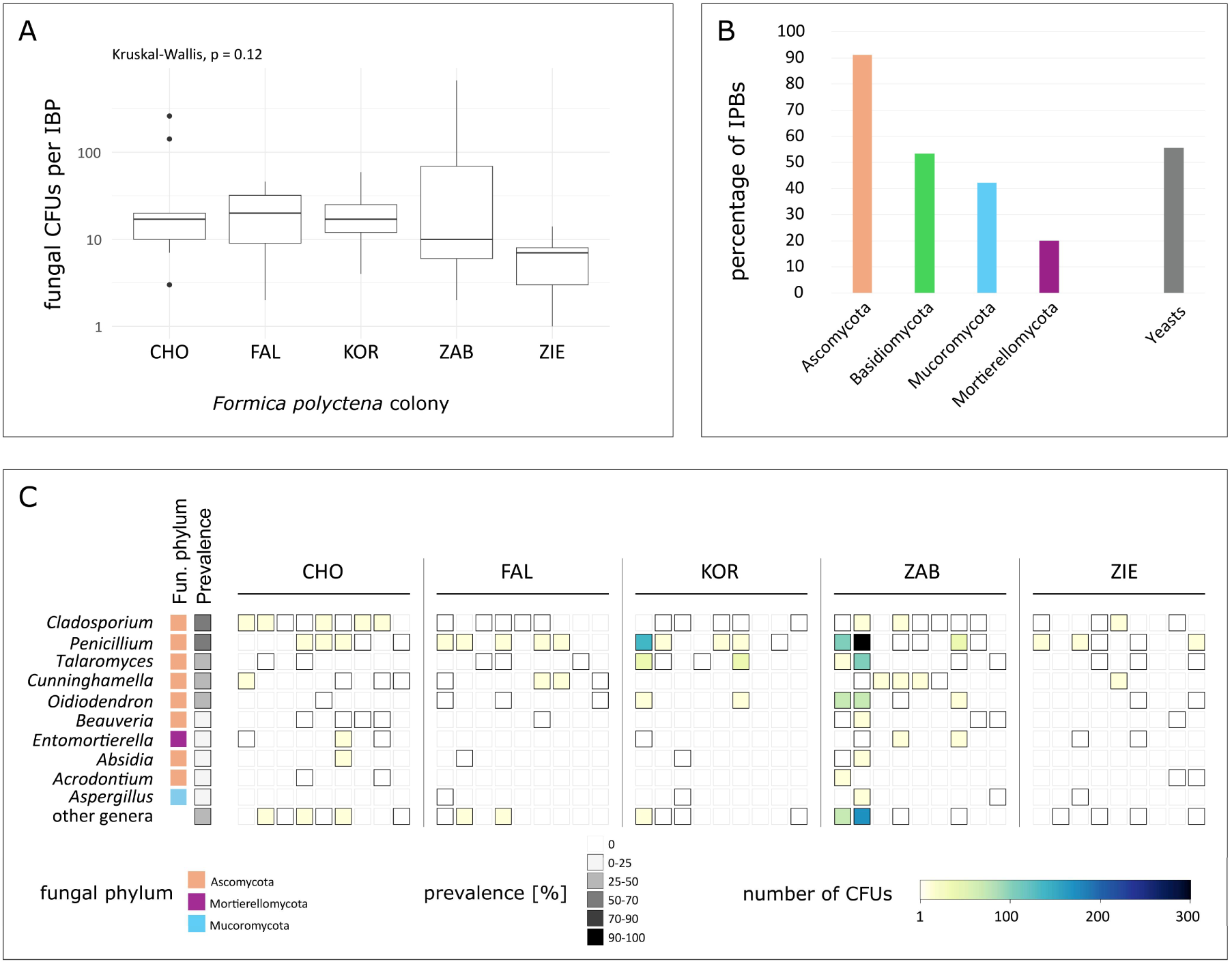
Diversity and abundance of fungal strains cultured from ants’ infrabuccal pockets (IBPs). **A.** The average abundance of fungal strains (CFU) per IBP grouped by ant colonies. **B.** The prevalence of filamentous fungi belonging to different phyla and yeasts. Prevalence is understood as the percentage of IBPs in which a given taxon was recorded. **C.** Prevalence and abundance (CFU) of filamentous fungal genera per IBP. Category “other genera” contains all filamentous fungal genera which did not exceed 10% prevalence. In the graph: letters (KOR, FAL, CHO, ZAB, ZIE) represent different ant colonies.

The cultured filamentous fungal colonies represented four phyla: *Ascomycota*, which grew from 91% of IBPs; *Basidiomycota*, which grew from 53%; *Mucoromycota*, from 42%; and *Mortierellomycota*, from 20%. Yeasts, defined here as fungal strains growing in a single-celled, non-filamentous form, grew from 55% of the IBPs (Fig. 3B). Based on colony morphology, we classified sporulating, non-yeast strains into 34 genera (Supplementary Table ST.4). Basidiomycetes as non-sporulating mycelia were excluded from further analysis. The most widespread genera were *Cladosporium*, *Penicillium*, *Talaromyces*, *Cunninghamella*, and *Oidiodendron*, isolated from 65%, 58%, 36%, 31%, and 27% of the IBPs, respectively. We observed representatives of *Cladosporium*, *Penicillium*, *Talaromyces*, and *Oidiodendron* in ants from every colony. *Penicillium* was the most abundant genus, averaging 16 CFUs per IBP (range: 0–295) (Fig. 3C).

We obtained 176 Sanger sequences (160 ITS and 16 LSU) from 185 strains representing 119 morphotypes. We used sequence data to identify 74 taxonomic units below the genus level (Supplementary Table ST.4). The most common isolates were representatives of the *Cladosporium cladosporioides* complex SH0962330.10FU, which grew from 47% of the pellets. We also isolated *Penicillium* SH0939727.10FU, the *Cladosporium herbarum* complex SH0962330.10FU, and *Talaromyces* SH0973349.10FU from 40%, 38%, and 31% of the pellets, respectively. Additionally, for 11 morphotypes, sequences obtained from representative strains did not closely match any entries in the NCBI database (ITS sequence identity <97%). Based on two such strains, we recently established a new fungal genus, *Formicomyces* (Siedlecki et al. 2023). Detailed information on the number of CFUs of each taxonomic unit in each IBP is provided in Supplementary Table ST.5.

### 3.3. Fungal diversity assessed using multi-marker DNA metabarcoding

For 57 out of 60 processed infrabuccal pellets, we obtained >1000 reads per sample, classified as non-contaminants (based on comparison with negative control data), for each of the three targeted regions: ITS2, ITS1, and 18S rRNA. Using these non-contaminant data, we reconstructed a total of 4214, 3212, and 3066 unique genotypes (ASVs) for the ITS2, ITS1, and 18S rRNA marker regions, respectively. Of these, 333, 346, and 225 ASVs, respectively, reached at least 1% relative abundance in at least one sample. The average number of ASVs per pellet varied across markers: 138 (range: 23–357) for ITS1, 173 (range: 30–465) for ITS2, and 118 (range: 24–326) for 18S rRNA.

For all molecular markers, more than 90% of ASVs were assigned to the phylum level with a confidence of ≥0.9. However, the success of assignments at lower taxonomic ranks varied among markers. For 18S rRNA ASVs, we confidently (i.e., with confidence ≥ 0.9) assigned 45% to the order level and 19% to the genus level. In contrast, for ITS1 and ITS2, 81% and 86% of ASVs were assigned to the order level, and 68% and 71% to the genus level, respectively, at the same confidence threshold (see Appendix A.10).

From 57 IBPs, we identified fungi from 11 phyla. *Ascomycota* were the most abundant, representing 81%, 66%, and 73% of the average relative abundance per IBP in the ITS2, ITS1, and 18S datasets, respectively. *Basidiomycota* constituted 12%, 25%, and 12% of the average relative abundance per IBP in the ITS2, ITS1, and 18S datasets, respectively. *Mortierellomycota* and *Mucoromycota* did not exceed 10% of the average relative abundance per IBP in any dataset. All other phyla comprised less than 1% of the average relative abundance per IBP (Fig. 4A). Across the three marker regions, we detected a total of 578 fungal genera, but with substantial differences among regions. For each region, we detected unique genera: ITS2 – 109 (19%), ITS1 – 71 (12%), and 18S – 38 (7% of all recorded genera). Only 69 genera (12%) were captured by all three marker regions, whereas approximately half of all reconstructed genus-level fungal diversity was captured by both ITS1 and ITS2 marker regions (Fig. 4B).

**Figure 4.**
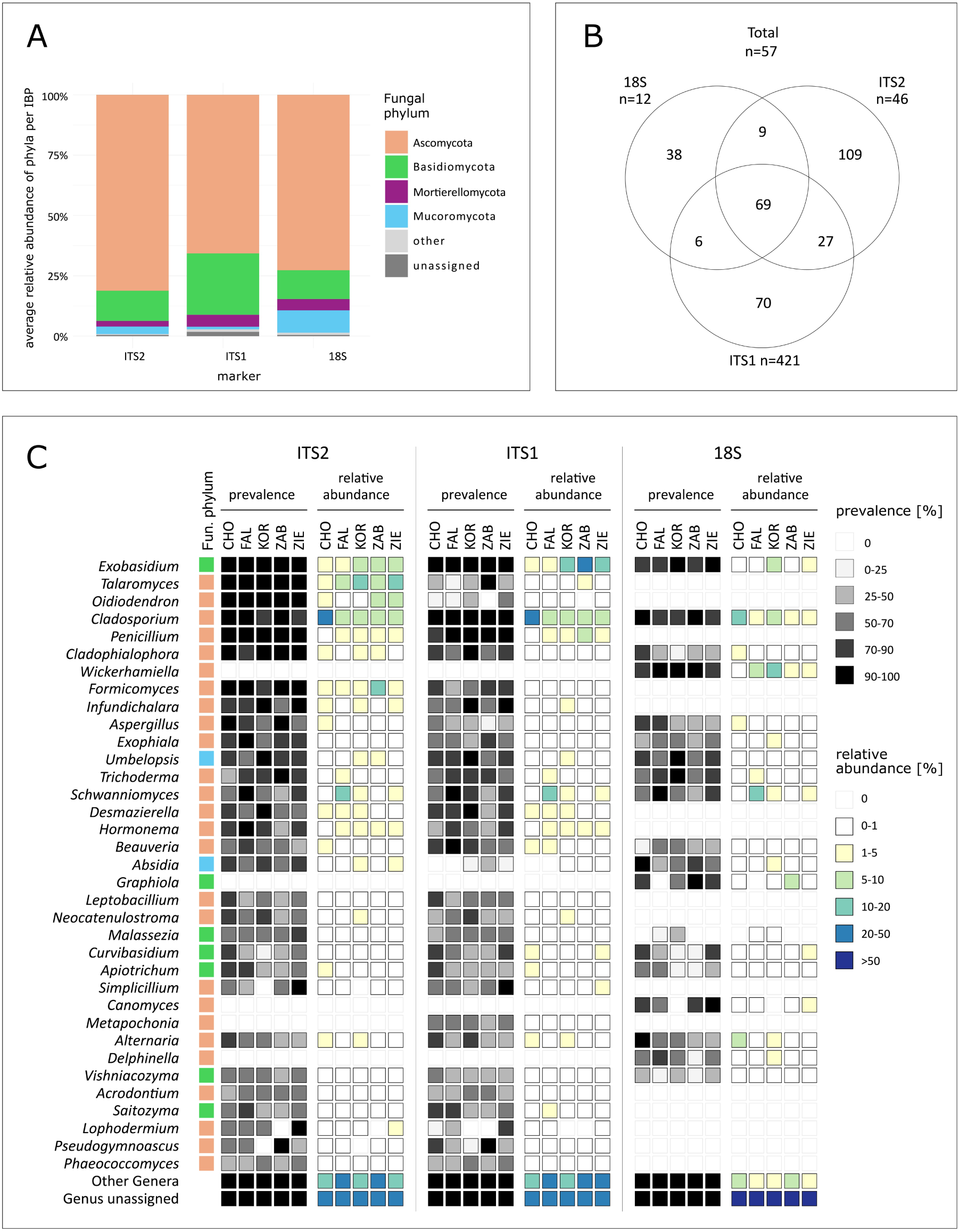
Fungal diversity and distribution in IBPs observed by multi-target (ITS2, ITS1, 18S rRNA) amplicon sequencing approach. **A**. The average relative abundance of fungal phyla per IBP, based on three marker regions. Category “Other” groups fungal phyla with average relative abundance <1%. **B**. An overlap among the three marker regions in lists of fungal genera identified. **C**. The distribution and average relative abundance per IBP of the most prevalent fungal genera across ant colonies. Prevalence is the percentage of IBPs in which a given genus was recorded. Category “Other Genera” contains all fungal genera that did not reach 50% total prevalence noted for any of the marker genes.

Fungi from 35 genera were detected in at least 50% of IBPs. The most prevalent genera, recorded in more than 90% of IBPs, included *Exobasidium*, *Talaromyces*, *Oidiodendron*, *Cladosporium*, *Penicillium*, *Cladophialophora*, *Wickerhamiella*, and *Formicomyces*. Additionally, *Infundichalara*, *Aspergillus*, *Exophiala*, *Umbelopsis*, *Trichoderma*, *Schwanniomyces*, and *Desmazierella* were present in more than 70% of IBPs (Fig. 4C). The most abundant genera were *Exobasidium* (13.0% average relative abundance in the ITS1 dataset, range: 0.7–64.0%), *Cladosporium* (10.5% average relative abundance in the ITS2 dataset, range: 0–50.0%), *Talaromyces* (8.5% average relative abundance in the ITS2 dataset, range: 0.1–43.0%), and *Wickerhamiella* (5.5% average relative abundance in the 18S dataset, range: 0–34%). Other genera did not exceed 5% in any dataset (Fig. 4C).

Most genera were relatively evenly distributed across ant colonies, but some fungi were substantially more common in certain colonies. For example, *Cladosporium* comprised, on average, 23.5% of ITS2 reads in ants from colony CHO, whereas in other colonies, its relative abundance was less than half of that, ranging between 6% and 10% (Fig. 4C). Similarly, based on ITS2, *Formicomyces* were more abundant in colony ZAB (10% of reads on average) than in other colonies (range: 2–4%), and *Schwanniomyces* were more abundant in colony FAL (12%) than in others (range: 0.1–4%). Furthermore, in the ITS1 dataset, *Exobasidium* reads were on average more abundant in colonies ZAB (22%), KOR (20%), and ZIE (14%) than in CHO (5%) and FAL (4%). Finally, in the 18S dataset, *Wickerhamiella* reads were on average more abundant in colony KOR (16%) than in others (range: 1–7%) (Fig. 4C).

Additionally, across the three marker regions, a large proportion of ASVs, together comprising a significant fraction of the reconstructed communities and sometimes broadly distributed, lacked genus-level assignments. In the most informative ITS2 dataset, the most prevalent such ASVs: ASV8 (*Capnodiales*), ASV27 (*Dothideomycetes*), and ASV31 (*Capnodiales*), occurred in 84%, 77%, and 75% of IBPs, respectively (see Appendix A.11). The distribution and relative abundance of each ASV across individuals for all marker genes are provided in Supplementary Tables ST.7-9.

### 3.4. Comparison of methods to study fungal diversity

The numbers of fungal genera obtained from IBPs using three independent approaches (microscopy, culturing, and multi-marker metabarcoding) differed. Multi-marker metabarcoding allowed us to detect the highest number of genera (578 in 57 IBPs), followed by culturing, which resulted in 34 genera in 45 IBPs, with the caveat that the taxonomic diversity of yeasts and sterile strains was not analyzed. Using direct microscopy, we were not able to achieve genus-level identifications as the distinguished morphotypes represented taxonomically collective categories encompassing a diverse range of fungal taxa. However, only direct microscopy provided direct evidence of fungal presence in the IBPs and allowed us to observe diverse life stages of fungi within the organ (fungal hyphae, spores, and single cells) (Fig. 2).

Detection of the seven most prevalent fungal morphotypes identified by direct microscopy, using culturing and multi-marker metabarcoding, differed between the methods. In the case of metabarcoding, for each of the seven morphotypes, we found at least one highly prevalent, morphologically matching fungal genus. In the case of fungal culturing, genera corresponding to microscopic morphotypes were in most cases isolated less frequently than suggested by microscopy data. Only yeasts were noted similarly prevalent between microscopy and culturing, with the caveat of limited taxonomic resolution in both approaches (Fig. 5).

**Figure 5.**
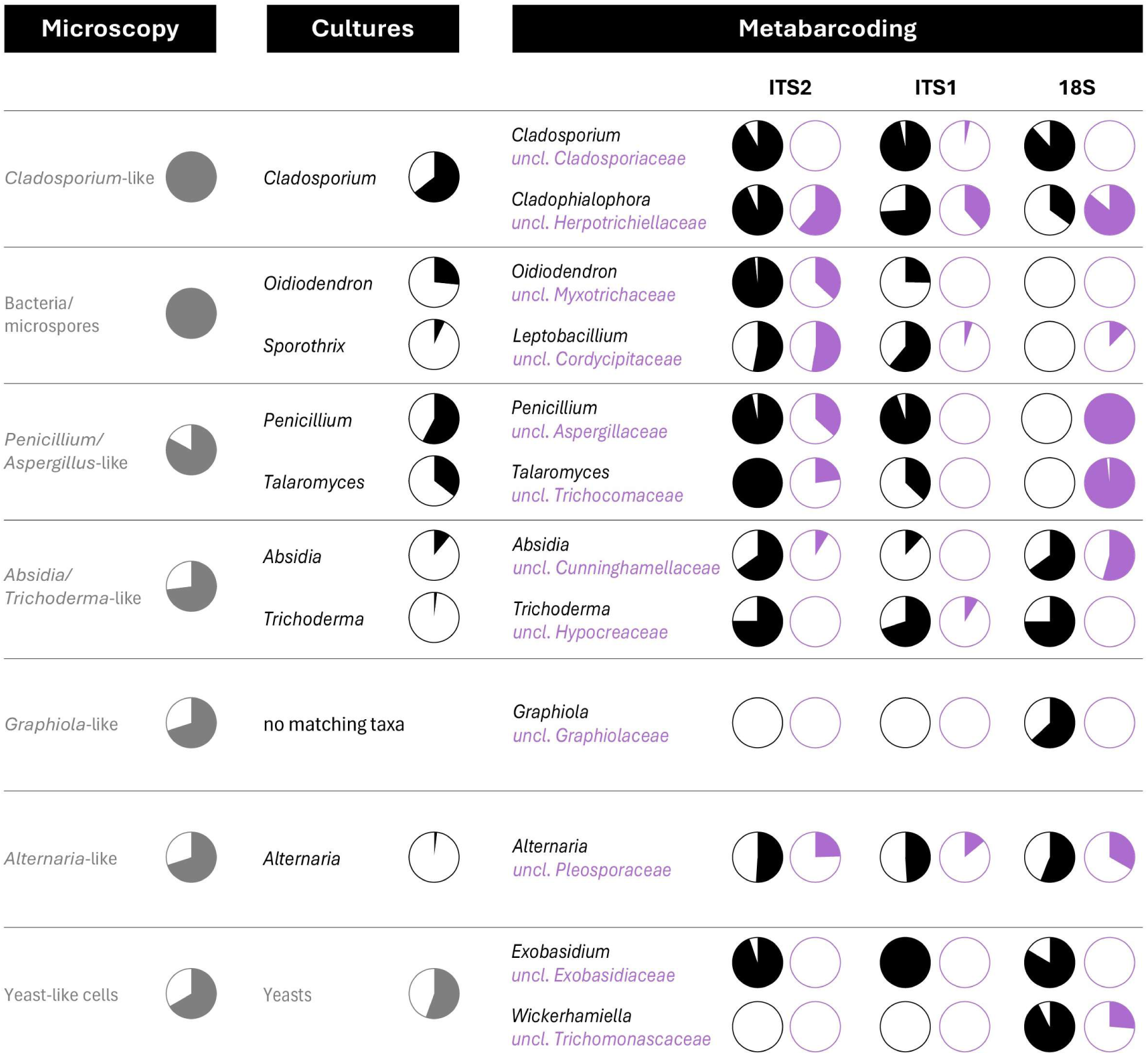
Detection of the most prevalent fungal morphotypes noted in direct microscopy through the use of culturing and multi-marker metabarcoding approaches. Pie charts show a percentage of analyzed IBPs with the taxa present. For Cultures and Metabarcoding, up to two most prevalent taxa representing each morphotype detected by direct microscopy are shown. Morphologically identified bins that group distinct fungal taxa are shown in grey. For metabarcoding, unclassified genera in the same family as the representative genus are shown in purple.

Most of the 159 unique ITS sequences from cultured strains representing 71 taxonomic units closely matched sequences present in the ITS2 (71%) and ITS1 (56%) metabarcoding datasets (assuming >99% identity and >75% overlap of the metabarcoding ASV with the reference) (see Appendix A.12). For the majority of these sequences, genus-level taxonomic assignment based on shorter metabarcoding sequences were consistent with those based on the full ITS region (85% for ITS2, 82% for ITS1). In the remaining cases, ASVs were classified at a higher taxonomic level than the corresponding Sanger sequences (see Appendix A.12). For seven taxonomic units detected via culturing, we did not find any matching sequences in either ITS metabarcoding dataset (assuming identity >95% and >50% overlap). While four of these taxa were isolated from a single IBP, three taxa: *Penicillium* SH0871105.10FU, *Penicillium* SH0940202.10FU, *Penicillium* SH0940189.10FU were isolated more prevalently, respectively 20%, 15.56%, 4.44% of IBPs. On the other hand, metabarcoding data included 547 genera that were not detected in the culturing data, although we presume that some strains representing these taxa grew in “yeast” or “sterile *Basidiomycota*” form that we did not attempt to classify in the culturing experiment.

### 3.5. Bacterial diversity assessed using 16S rRNA metabarcoding

For 59 out of 60 samples, we obtained more than 1000 16S rRNA reads, which were classified as non-contaminants based on comparisons with negative controls (average: 34,150, range: 2,996–90,751). These non-contaminant reads represented 2,497 ASVs, with an average of 82 ASVs per IBP (range: 6–265). These ASVs clustered into 831 97% OTUs, representing 144 families across 37 classes.

Bacteria from four classes: *Bacilli*, *Alphaproteobacteria*, *Gammaproteobacteria*, and *Actinobacteria* comprised, respectively, 54%, 32%, 11%, and 2% of reads per IBP on average, with none of the other classes exceeding 1% (Fig. 6A). Three bacterial OTUs from these classes were present in at least 90% of IBPs - otu2 (*Fructilactobacillus*), otu4 (*Oecophyllibacter*), and otu10 (*Wolbachia*). Four additional OTUs were present in at least 70% of IBPs: otu6 (*Acetobacteraceae*), otu9 (*Serratia*), otu33 (*Burkholderia*), and otu14 (*Lactobacillus*) (Fig. 6B). The same OTUs were also among the most abundant in the dataset. In particular, otu2 (*Fructilactobacillus*) constituted 52% (range: 2–93%) of reads per IBP on average, followed by otu4 (*Oecophyllibacter*) and otu6 (*Acetobacteraceae*), representing 11% (range: 0–49%) and 10% (range: 0–63%) of reads per IBP on average (Fig. 6B). Some other bacteria were less evenly distributed among colonies. For example, otu7 (*Acetobacteraceae*) and otu8 (*Serratia*) were both common and abundant in colony ZIE, but much less so in others. The distribution and relative abundance of each 16S rRNA ASV and OTU across individuals are provided in Supplementary Tables ST.10-11.

**Figure 6.**
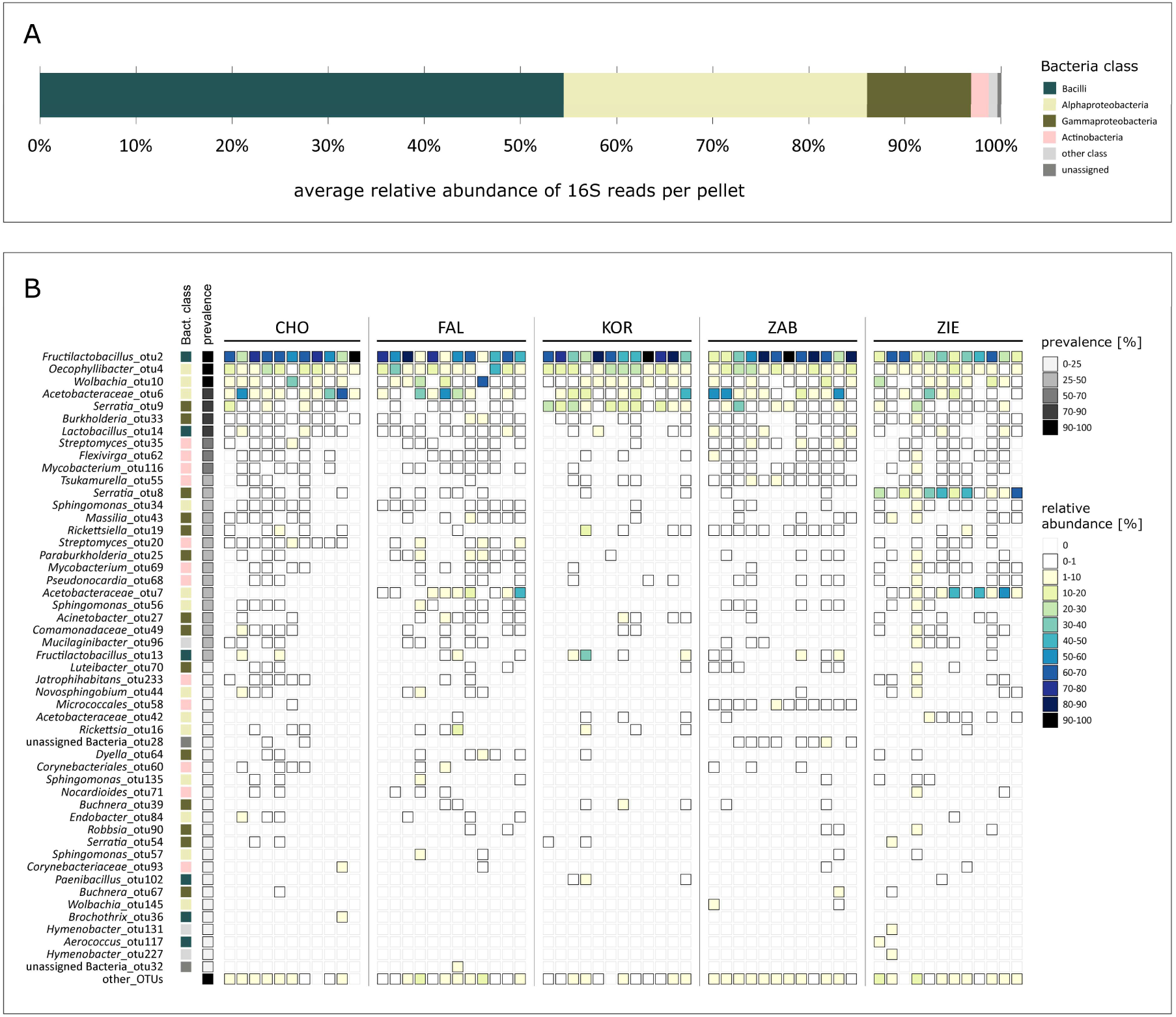
The distribution of bacterial OTUs across IBPs of *F. polyctena* worker individuals from five colonies, based on amplicon sequencing data for the V4 region of the 16S rRNA gene. **A** The average relative abundance of bacterial classes in infrabuccal pellets. Category “other class” contains those bacterial classes which did not reach 1% of average relative abundance. **B** The distribution and average relative abundance of the bacterial OTU across ant colonies. Category “other OTUs” contains all OTUs that did not exceed 1% of relative abundance.

### 3.6. General diversity patterns in fungal and bacterial communities

To assess general diversity patterns, we used the fungal ITS2 dataset, which had the highest number of ASVs and the highest confidence of assignment to the genus level, alongside the bacterial 16S rRNA dataset. While ant colonies did not differ significantly in the number of fungal ITS2 ASVs (Kruskal-Wallis test, p = 0.54, Fig. 7A), they did differ significantly in the number of bacterial ASVs (Kruskal-Wallis χ² = 11.792, df = 4, p = 0.019, Fig. 7A). Significant differences in community composition among colonies were detected in both 16S rRNA and ITS2 datasets (PERMANOVA, 16S: R² = 0.1527, p = 0.001; ITS2: R² = 0.1552, p = 0.001; pairwise comparisons, p < 0.01). However, for 16S rRNA data, pairwise comparisons indicated significant differences only between some colonies: KOR vs. FAL (p = 0.02), KOR vs. ZIE (p = 0.01), and FAL vs. ZIE (p = 0.04). In contrast, for ITS2, all pairwise comparisons revealed significant differences (p < 0.01) (Fig. 7B).

**Figure 7.**
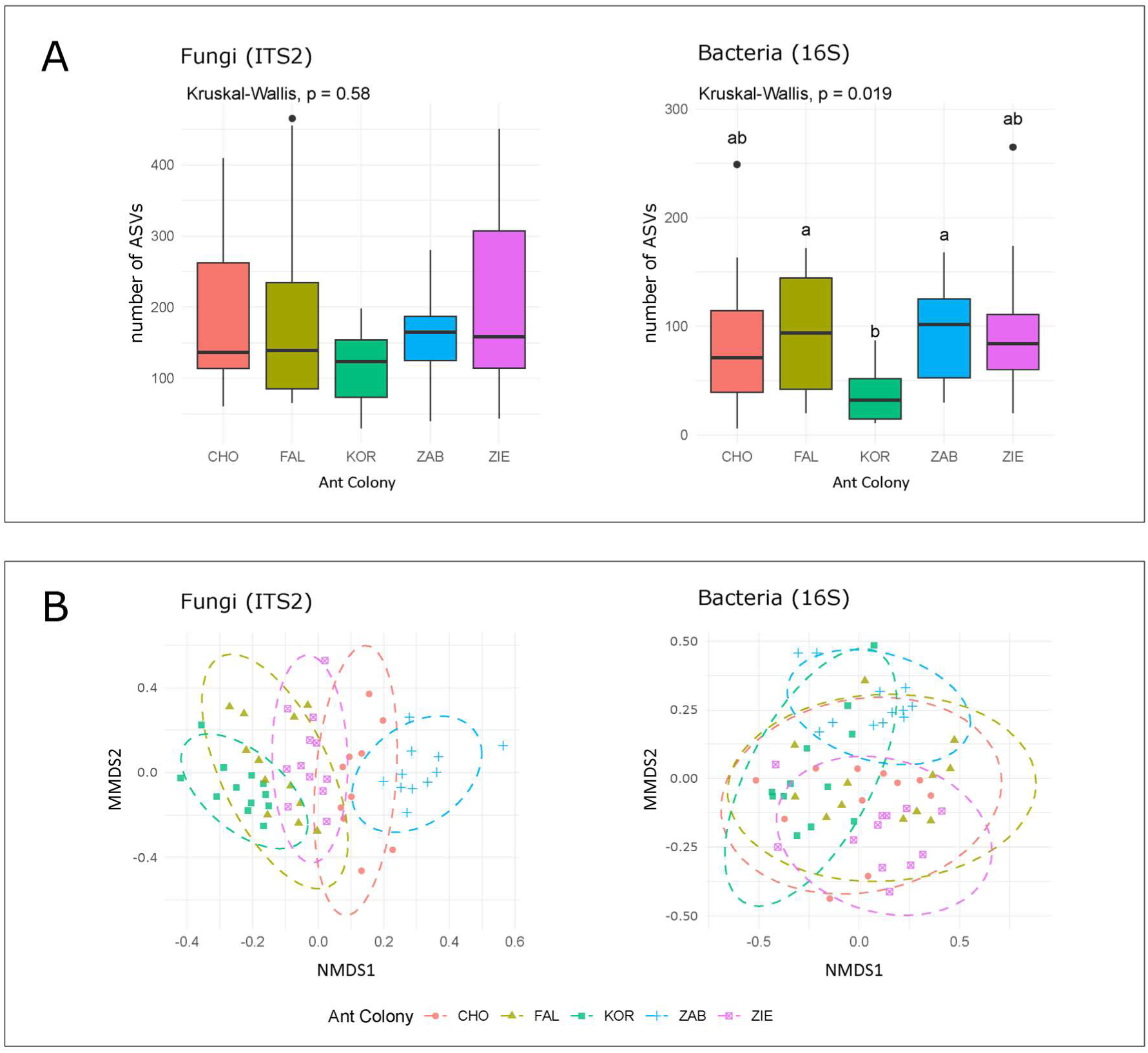
Determination of general diversity patterns (**A-B**) of bacterial (16S) and fungal (ITS2) communities. **A**. Boxplots showing ASV counts for the two communities grouped by ant colonies. The letters (a, b) above the boxplots represent the significant differences in the number of ASVs between colonies. **B**. Non-metric multidimensional scaling (NMDS) ordinations of the composition of the two communities based on Bray–Curtis distances. Points represent individual samples, and ellipses depict 95% confidence intervals for group centroids. Colors and shapes correspond to different colonies.

**Figure 8.**
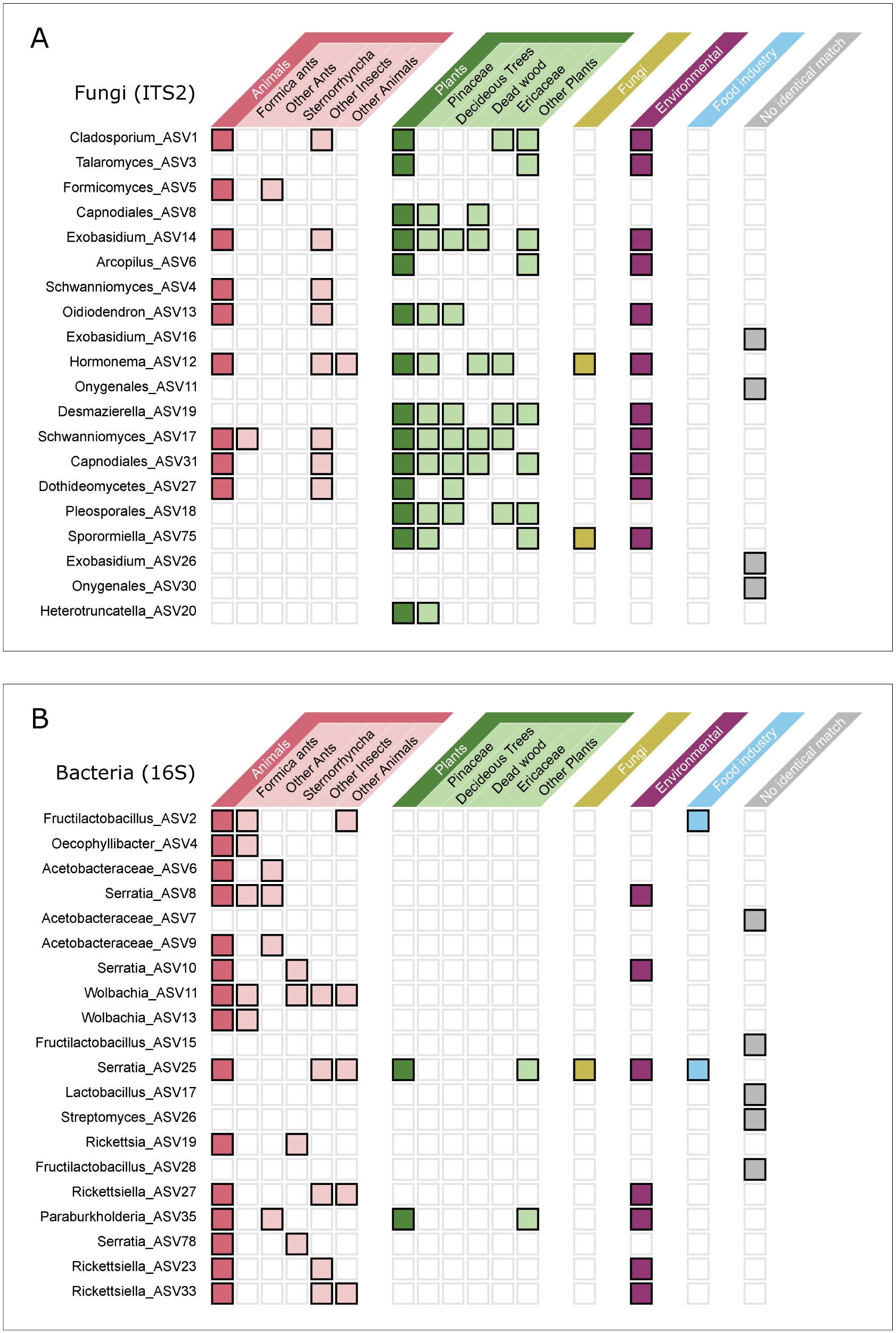
The ecological distribution of the most abundant fungal (**A**) and bacterial (**B**) ASVs from ant infrabuccal pockets, determined based on the isolation source of 10 matching sequences per ASV available in the NCBI database. Tiles with bolded edges indicate matching of ASV with reference genotypes, and color of the tile refers to the genotypes isolation source. The “environmental” category refers to sequences isolated from air, water, or soil samples.

### 3.6. Ecological specificity of fungal and bacterial communities

To assess ecological specificity, we determined the environmental distribution of the core ITS2 and 16S ASVs based on metadata associated with identical sequences in the NCBI core nucleotide database (Supplementary Table ST.12). Of the 20 most abundant fungal ASVs in our dataset, only one has been previously reported from *Formica* ants, and one from other ants. In total, 45% of these ASVs were previously reported from insects. At the same time, 70% of these fungal ASVs have been reported from plants, most often conifers, deciduous plants, heathers (*Ericaceae*), and dead wood. 55% of these fungal ASVs have also been reported from the environmental samples - soil, water, or air (Fig. 7C). In comparison, for the bacterial community, 75% of the 20 core 16S rRNA ASVs matched sequences previously isolated from animals. Of these, 25% of ASVs represented genotypes matching *Formica* ant-isolated sequences, 20% have been found in other ants, 20% in sap-sucking insects (*Sternorrhyncha*), and 25% in other insects. Only 35% of these core bacterial ASVs matched environmental sequences, and 10% sequences were isolated from plants (Fig. 7C). For 25% of bacterial and 20% of fungal ASVs no identical sequences were found in the reference database.

## 4. Discussion

### 4.1. The value of implementing a multi-method approach to study fungal communities

The use of alternative methods for the study of ant-associated fungal communities provided us with very different perspectives and a more comprehensive overview of the IBP mycobiome. Fungal diversity studies have long relied on culturing and direct microscopy approaches, despite the limitations of these classical methods: their time-consuming nature, the need for taxonomic expertise, and the non-cultivability of many fungal taxa (Dziurzynski et al. 2023). Metabarcoding overcomes some of these limitations (Lücking et al. 2020; Xu, 2016), but brings several of its own. Combining classical methods with metabarcoding enabled their side-by-side comparison in an ecological context.

The main limitation of the direct microscopy approach was its low taxonomic resolution, forcing the assignment of morphotypes to collective categories, sometimes grouping distantly related taxa, e.g., representatives of *Cladosporium* (*Dothideomycetes*) and *Cladophialophora* (*Eurotiomycetes*) (Ho et al. 1999). Since the material collected in the IBP is compacted into a pellet, the observed fungi lacked key morphological characteristics, such as the mode of conidiogenesis, necessary for precise taxonomic identification (Seifert et al. 2011; Skirgiełło et al. 1979). On the other hand, unlike the other methods, direct microscopy provided direct evidence of diverse fungal structures present within the material.

In culturing, morphological features combined with DNA barcodes enabled the identification of most colonies (80%) at the genus level, but with important exceptions. Specifically, morphological identification was not possible for sterile cultures (3% of fungal CFUs) which lacked reproductive structures crucial for morphological classification (e.g., Seifert et al. 2011), and for yeasts (10% of fungal CFUs), which require selective media and physiological comparisons as part of the identification process (Kurtzman et al. 2011). Another challenge is the method’s bias toward fast-growing and abundantly spore-producing species (Quan et al. 2019; Warcup, 1950). Indeed, in our study, slow-growing *Chaetothyriales* (Quan et al. 2020), common in metabarcoding data, were rarely or not at all isolated through culturing. Despite these limitations, culturing was the only method enabling partial verification of whether the fungi observed in the IBP were viable and the acquisition of stable cultures necessary for the formal description of new taxa (Aime et al. 2021), expansion of reference databases, metabolite profiling, or experiments.

In turn, the outcomes of metabarcoding depended on the targeted region. ITS2 and ITS1 markers provided high taxonomic resolution, with ∼70% of ASVs confidently identified to the genus level, which is consistent with previous findings (e.g., Tedersoo et al. 2015). However, the use of the conserved 18S rRNA marker, which offered much lower taxonomic resolution, still allowed us to detect genera commonly present in the IBP but lacking from ITS2 or ITS1 datasets, such as *Canomyces*, *Delphinella*, *Graphiola*, and *Wickerhamiella*. This finding aligns with the known limitations of ITS regions and supports the general recommendation to use multi-marker DNA metabarcoding for the characterization of fungal communities (Lücking et al. 2020; Płoszka et al. 2025; Xu et al. 2016). Moreover, a well-known limitation of molecular identification in metabarcoding is its strong dependence on the completeness of the reference database (Lücking et al. 2020). As of February 2025, the UNITE reference database contained approximately 3.8M ITS sequences clustered into ∼2.4M species hypotheses (SHs), most of which lack links to recognized fungal species (Abarenkov et al. 2024). The implementation of culturing in our study enabled the isolation of two unknown fungal strains, based on which a new genus and species - *Formicomyces microglobosus* gen. et sp. nov. - were designated (Siedlecki et al. 2023). This allowed us to match ITS2 Zotu5, prevalent in the IBP, to the *Formicomyces* genus.

Overall, results obtained using different methods are complementary and jointly provide better insights into the biology of ant-fungal interactions. For example, while *Trichoderma*-like spores were observed in 73% of IBPs and ITS2 ASVs assigned to *Trichoderma* were detected in 74% of IBPs, the genus was successfully cultured from only 2% of the IBPs. These findings suggest that *Trichoderma* spores become inactivated while stored in the IBP, as species of this genus are generally culturable (Harman and Kubicek, 2002), and strains of *Trichoderma* commonly grew readily under the same culturing conditions when *F. polyctena* cadavers were used as a source (Siedlecki et al. 2021). Two other easily culturable genera, entomopathogenic *Beauveria* and *Simplicillium* (Chen et al. 2019; Rehner et al. 2011), were also cultured less frequently from the pellets (22% and 4%, respectively), than they were observed using the ITS2 metabarcoding approach (60% and 50%, respectively). The inactivation of fungal spores in IBPs has been previously reported for *Atta* and *Lasius* ants (Fernández-Marín et al. 2006; Tragust et al. 2013), and our study suggests that *F. polyctena* should be added to the list. This finding supports the view of the IBP as an organ involved in ant immunity (Fernández-Marín et al. 2006; Tragust et al. 2013).

For bacteria, the majority of which remain unculturable under standard conditions (Steen et al., 2019), 16S rRNA amplicon sequencing has become the preferred method for assessing bacterial communities, despite its various caveats. However, culturing remains an important element of in-depth characterization of host-bacteria associations, essential for reconstructing interaction mechanisms (Akami et al., 2019; Kwong et al., 2017).

### 4.2. Who are the microorganisms?

Ant clades differ in the abundance and specificity of their associated microbes, ranging from specialized symbionts that are reliably transmitted transovarially or socially (Hu and Moreau, 2025; Degnan et al. 2004; Ramalho et al. 2018) to low-abundance and inconsistent associations (Hammer et al. 2019; Sanders et al. 2017). Our findings indicate that the bacterial community within the IBP is largely consistent across colonies, aligning with patterns observed in ant clades harboring specialized microbes. Moreover, bacterial communities found in the IBP of *F. polyctena* primarily consist of taxa previously reported from the mound-building *Formica* species, other ants and insects, including *Lactobacillaceae*, *Wolbachia*, and *Acetobacteraceae* (Jackson et al. 2023; Sinotte et al. 2024; Zheng et al. 2022). However, a growing number of studies suggest that different parts of the ant digestive tract are inhabited by distinct bacterial communities (Flynn et al. 2021; Lanan et al. 2016). This is likely the case here: the high relative abundance of *Fructilactobacillus* (*Lactobacillaceae)* in *F. polyctena* IBP, consistent with previous reports of *Lactobacillaceae* high abundance in the IBP and crops of three other *Formica* species (Zheng et al. 2022), suggest specialization to thrive specifically in the initial parts of these ants’ digestive tract. This specialization might be driven by *Lactobacillacae* being tolerant of lower pH (Ng et al. 2023) in the ant infrabuccal pockets (Tragust et al. 2020). On the other hand, representatives of *Wolbachia* are endosymbionts known to reside within host insect tissues and hemolymph (Saridaki and Bourtzis, 2010), which explains their consistent presence, but relatively low abundance in IBP compared to previous studies that used homogenized individuals (Jackson et al. 2023; Kaczmarczyk-Ziemba et al. 2020; Sinotte et al. 2024). In turn, the consistent detection in *F. polyctena* IBP of *Oecophyllibacter* (*Acetobacteraceae*), originally cultured from SE Asian weaver ant *Oecophylla smaragdina* (Chua et al. 2020), suggests a broader association of this bacterial clade with ants.

Regarding fungi, some commonly detected taxa have also been reported in previous culture-based studies of the *Formica* mycobiome. These include species of *Wickerhamiella* and *Schwanniomyces*, reported from red wood ants and their nest material (Golubev and Bab’eva, 1972; Golubev and Bab’eva, 1977; Maksimova et al. 2016), as well as from other habitats (e.g., flowers, soil, human samples, and alcoholic beverages) (de Vega et al. 2017; Lachance et al. 1998; Suzuki and Kurtzman, 2011; Takei et al. 2024). Moreover, representatives of *Penicillium*, which were commonly detected in our study, have previously been abundantly isolated from the cadavers of *F. polyctena* (Siedlecki et al. 2021). Finally, a notable finding is the high prevalence of *Formicomyces* (Trichomeriaceae) in IBPs, as representatives of this genus have previously only been reported as *Chaetothyriales* sp. 1, from the carton nests of *Lasius fuliginosus* and *Lasius umbratus* (Schlick-Steiner et al. 2008). While representatives of the closely related genus *Trichomerium* (*Trichomeriaceae*) have been shown to colonize the carton nests of tropical and subtropical ants (Quan et al. 2020), our findings support the hypothesis of an association between *Formicomyces* and temperate ants (Siedlecki et al. 2023).

The IBP microbial community composition is suggestive of strong interaction between ants and their tended sap-sucking insects (*Sternorrhyncha*). The first argument is the high prevalence of *Serratia*, commonly associated with these sap-sucking insects (Pons et al. 2019) and detected in the guts of ants tending aphid colonies infected with this bacterium (Renoz et al. 2019). Another argument is the high prevalence of sooty molds (e.g., *Alternaria*, *Cladosporium*, *Chaetothyriales*, *Capnodiales* s. str.), known to grow on sugary exudates of plants and sap-sucking insects (Abdollahzadeh et al. 2020; Levetin and Dorsey, 2006; Magyar et al. 2023; Quan et al. 2020). We believe that sooty molds may be collected into the IBP during the ants’ aphid-tending activities. Given the growing evidence that some honeydew-associated taxa, such as *Cladosporium*, may be entomopathogenic to aphids (Nicoletti et al. 2024), the collection of potentially pathogenic spores into the IBP could play an important role in the ant-aphid mutualism.

However, many bacterial and fungal taxa found in the IBP have not previously been reported as ant- or insect-associated, but instead have much broader environmental distributions, suggesting their external origin and opportunistic nature. This is particularly evident for the fungal community, comprising multiple fungal genotypes previously reported from plants. Moreover, as most fungi inhabiting insect guts occur in a yeast-like form (Blackwell, 2017; Gibson and Hunter, 2010), the common presence of fungal hyphae and spores in pellets further supports this hypothesis. This is likely the case for representatives of ubiquitous, plant-associated, or saprotrophic fungal genera such as *Cladosporium* (Bensch et al. 2010), *Penicillium* (Srinivasan et al. 2020), *Oidiodendron* (Rice and Currah, 2005), and *Talaromyces* (Kharkwal et al. 2024). We also recorded fungal taxa specifically associated with plants prevalent in pine forests, including pine-associated representatives of *Desmazierella* (Martinović et al. 2016) and *Lophodermium* (Osono and Hirose, 2011), as well as representatives of *Exobasidium*, which are common endophytes and pathogens of heathers and berries (*Ericaceae*) (Begerow et al. 2002). These fungi may grow and sporulate on plant organic matter that forms the mound, and be collected by ants during nest cleaning or nestmate grooming. This hypothesis is supported by findings of Lindström et al. (2021), who identified representatives of *Cladosporium*, *Penicillium*, *Oidiodendron*, and *Lophodermium* in the mound material of *Formica exsecta*.

### 4.3. What do IBP-associated microbes teach us about ant biology?

Previous research has shown that stable, core gut microbial communities in ants are closely linked to food specialization and nutritional supplementation (Hu et al. 2018; Hu and Moreau, 2025). The consistent presence in the *F. polyctena* IBP of osmotolerant, sugar-associated taxa such as lactic acid bacteria, particularly *Fructilactobacillus* (Endo and Salminen, 2013; Filannino et al. 2019), acetic acid bacteria (*Acetobacteraceae*) (Crotti et al. 2010), and *Wickerhamiella* yeasts (Gonçalves et al. 2018), indicates that these microorganisms are likely associated with the ants’ sugar-rich diet and may participate in the digestion and fermentation of honeydew-derived carbohydrates, as suggested for other ant clades (Brown and Wernegreen, 2016; Hu et al. 2018; Ivens et al. 2018; Russell et al. 2009; Sinotte et al. 2024). The high relative abundance of *Fructilactobacillus* may be specifically linked to the fact that we collected ants during the summer, a peak season for aphid-tending (Domisch et al. 2009); such seasonal shifts and consistent changes in *Lactobacillaceae* abundance have been reported from this ant species before (Sinotte et al. 2024). At this point, we do not know how these sugar-associated taxa affect ant nutrition and performance.

We can also infer some aspects of ant biology from the absence of certain taxa. For example, basidiomycetous macromycetes, which serve as an important food source for many invertebrate species (Santamaria et al. 2023), including ants (Epps and Penick, 2018; Schultz, 2021), were not commonly present in the IBP of *F. polyctena*. Our results suggest that macrofungi are not often foraged by *F. polyctena* ants. However, since our sampling was conducted in August, the rare occurrence of basidiomycetous macromycetes may be due to unfavourable dry conditions for their fruiting.

As the majority of fungal taxa detected in the IBP of *F. polyctena* are saprotrophs and plant associates, and the IBP is known to be regularly emptied (Richter and Economo, 2023), our study raises questions about the role of ants in fungal dispersal on the forest floor. While ant-mutualistic fungi are known to be stored and transported in the IBP (Hölldobler and Wilson, 1990; Mayer et al. 2023), less is known about the dispersal of non-mutualistic fungi. It has been suggested that species of *Glomus* are dispersed by harvester ants (Allen et al. 1984; Friese and Allen, 1993) and that fungus-farming ants may aid in the dispersal of saprotrophic fungi by expelling infrabuccal pellets onto refuse piles or at a distance from their nests (Mueller et al. 2001). Our observations that some plant-associated fungal taxa remain viable in IBP and can be successfully cultured from the pellets, add to the evidence of red wood ants’ roles in structuring fungal-plant interactions in forests.

### 4.4. Conclusions

Our study revealed a diverse and abundant microbial community associated with the infrabuccal pockets of *Formica polyctena* ants. The use of parallel methods and a multi-marker approach provided a more comprehensive understanding of the IBP mycobiota. In general, for both bacteria and the fungi, amplicon sequencing has emerged as an effective method for studying of their communities across many samples, offering a good balance between depth, throughput, cost, and effort. Nevertheless, the alternative approaches can provide valuable complementary insights and will continue to play important roles in microbiome studies.

The mycobiota within ant IBP comprised fungal hyphae, spores, and yeast cells belonging to diverse genera, with no single highly dominant taxon. Concurrently, the bacterial community was primarily composed of representatives of *Bacilli* and *Alphaproteobacteria*, with *Fructilactobacillus* being particularly abundant. A comparison of fungal and bacterial communities revealed the greater stability and higher ecological specificity of the latter. The careful interpretation of these diversity patterns provides a fresh perspective on the role of the IBP and its microbiome in ant biology, and on ants as ecosystem engineers shaping temperate forests.

## 5. Acknowledgements

We are thankful to Maria Majchrowska for her help in laboratory work, and Natalia Ciastoń for her help in graphical design of figures.

## 6. Authors Contribution

I.S. conceptualized the study; I.S. with help of M.W, J.P, and P.Ł designed the study; I.S., M.Koc., I.B. contributed to sampling and sample preparation; I.B. and M.W. conducted direct microscopy part with help of I.S.; I.S. with help of Z.B. and M.Koc. conducted culturing combined with representative strains’ barcoding; M.B. with help of I.S. and Z.P. conducted laboratory part of DNA metabarcoding; I.S. with help of M.Kol., and P.Ł conducted bioinformatic analysis of DNA metabarcoding; M.Koc. with help of I.S. performed statistical analysis; I.S., M.W., P.Ł performed data interpretation; I.S., K.N., I.B., and M.Koc. performed data visualization, I.S. with help of M.W., P.Ł., M.Koc., M.Kol., wrote the original version of the manuscript; M.W., P.Ł., J.P. - reviewed the manuscript.

## 8. Data Availability

Colony metadata are listed in Supplementary Table ST.1. Visual representations of all morphotypes distinguished in the direct microscopy part of the study are provided in Appendix AF.2-3, and raw data underlying results of this part of the study are provided in Supplementary Tables ST.2-3. Representative voucher specimens obtained during the culturing part are stored in the General Herbarium, University of Warsaw [WA], and sequence data generated for these specimens are available in the GenBank database. All reference numbers and taxon identification details for the specimens are provided in Supplementary Table ST.4. Raw data underlying results of the culturing part of the study are provided in Supplementary Table ST.5. Raw amplicon sequencing data from the multi-marker metabarcoding part of the study have been deposited in the Sequence Read Archive (SRA) of the National Center for Biotechnology Information (NCBI) under BioProject Accession no. PRJNA1185939 and are listed in detail in Supplementary Table ST.6. Decontaminated datasets from the metabarcoding part, used for analysis and visualization, are provided in Supplementary Table ST.7-11. All Supplementary Tables are available at: https://doi.org/10.58132/ZNZUKB. Scripts used for the bioinformatics analysis of multi-marker amplicon sequencing are available at https://github.com/IgorSiedlecki/Microbial-diversity-of-Formica-polyctena-infrabuccal-pockets. The R script for statistical analysis is available at https://github.com/mjkochanowski/Formica_polyctena_infrabuccal_pocket_microbiome. All occurrence-related data are available at GBIF: https://doi.org/10.15468/4hrjky, https://doi.org/10.15468/73qmpz, https://doi.org/10.15468/gvvcm9, https://doi.org/10.15468/m8usmu, https://doi.org/10.15468/sdekzb.

## Appendix

## A.1. An isolation of infrabuccal pellet from *Formica polyctena* ant

**Figure A.1.**
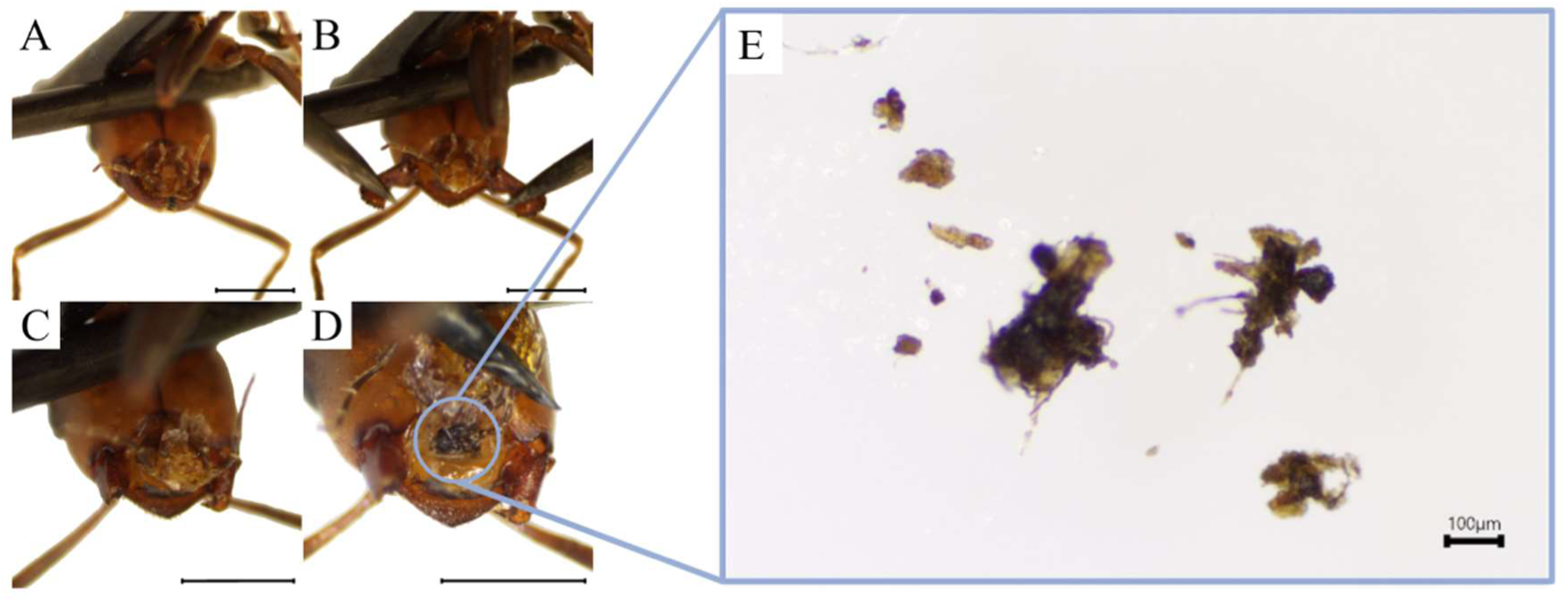
**A.** Each ant was placed dorsally on a dissecting sponge, with its head and body immobilised using 2–3 entomological pins. **B.** The mandibles were retracted with pins. **C.** Then severed using entomological scissors to facilitate further preparation. **D.** The lower lip was carefully pulled back using another pin to expose the infrabuccal pocket (highlighted by a blue circle). Laboratory equipment was flame-sterilised between individuals to avoid cross-contamination, and all procedures were performed on a clean bench. Scale bar in photos A-D: 1 mm. **E**. An isolated infrabuccal pellet placed on a microscope slide. The isolation was performed under a Nikon SMZ 745 T stereomicroscope.

## A.2. Biological diversity within infrabuccal pellets of *Formica polyctena* ants

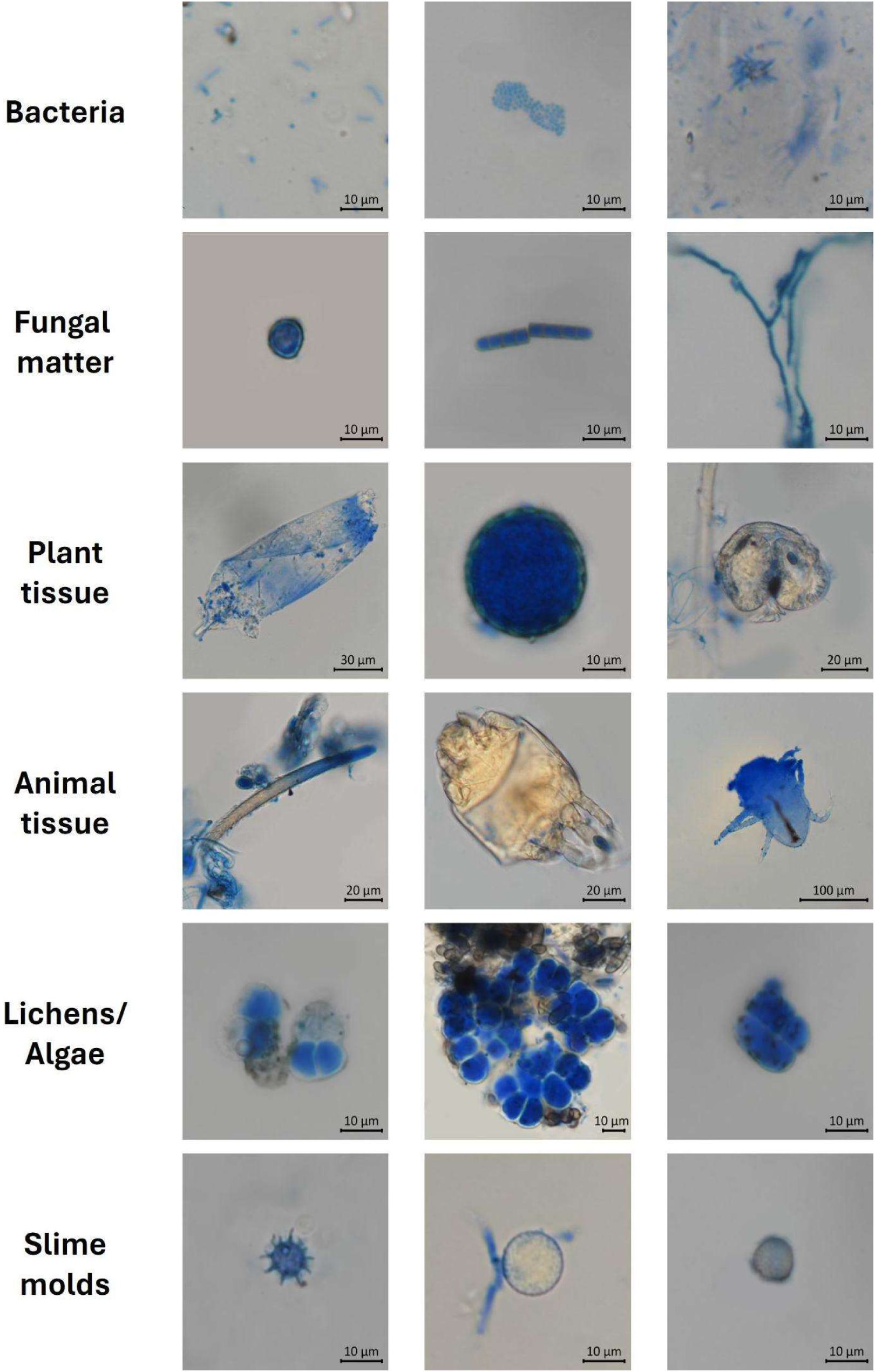

## A.3. Distinguished fungal morphotypes found in infrabuccal pellets (IBPs) of *Formica polyctena* ants

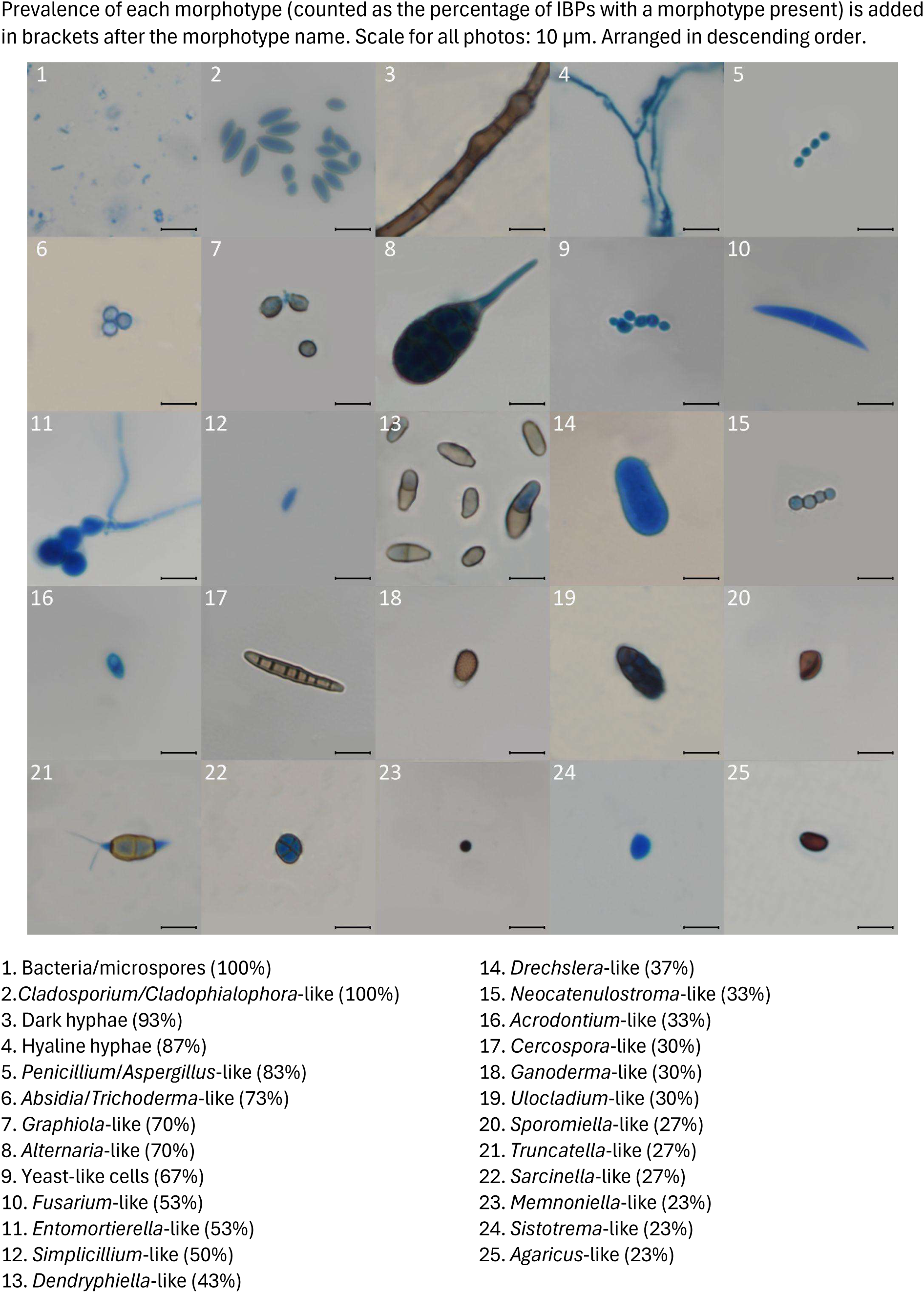
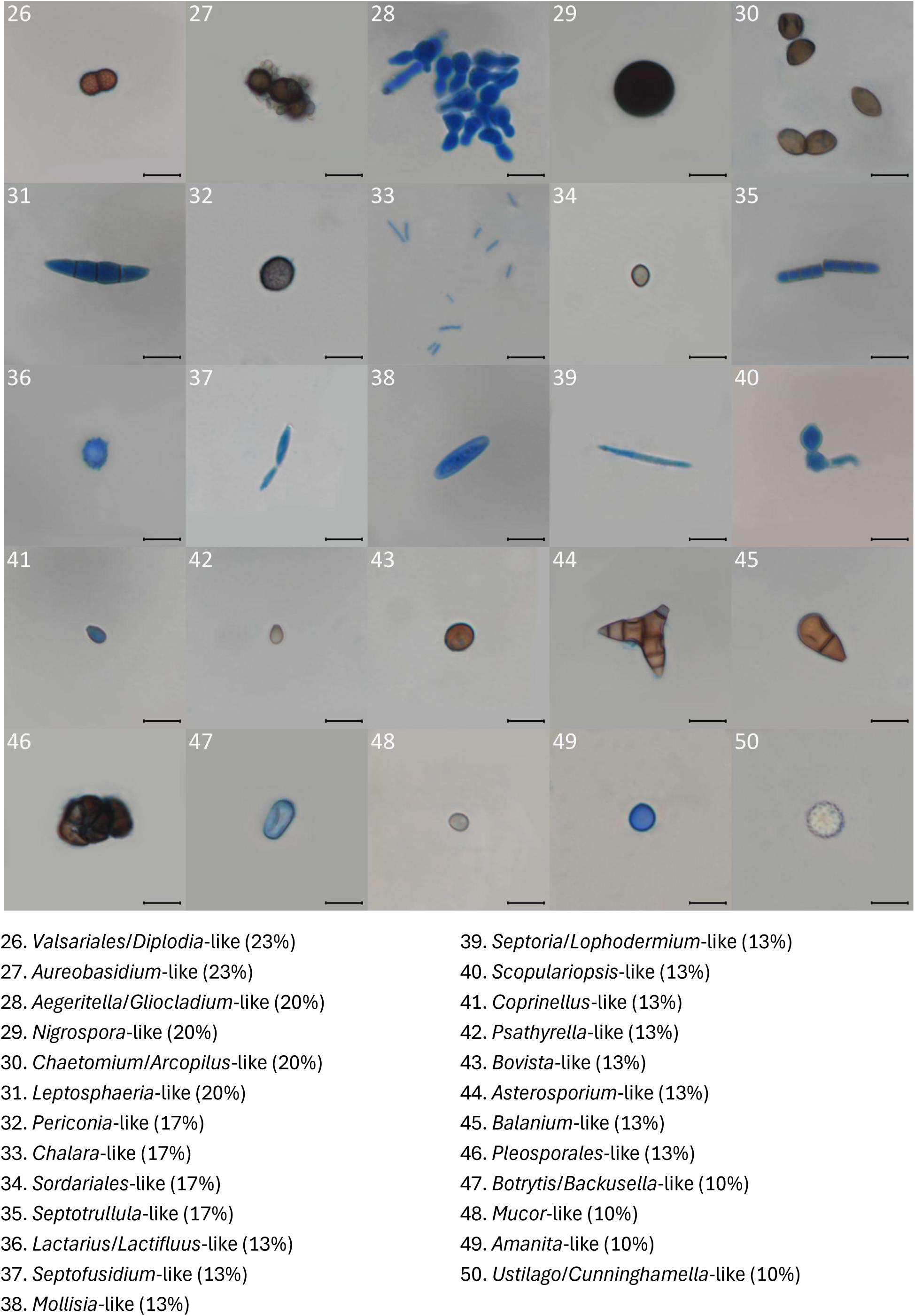
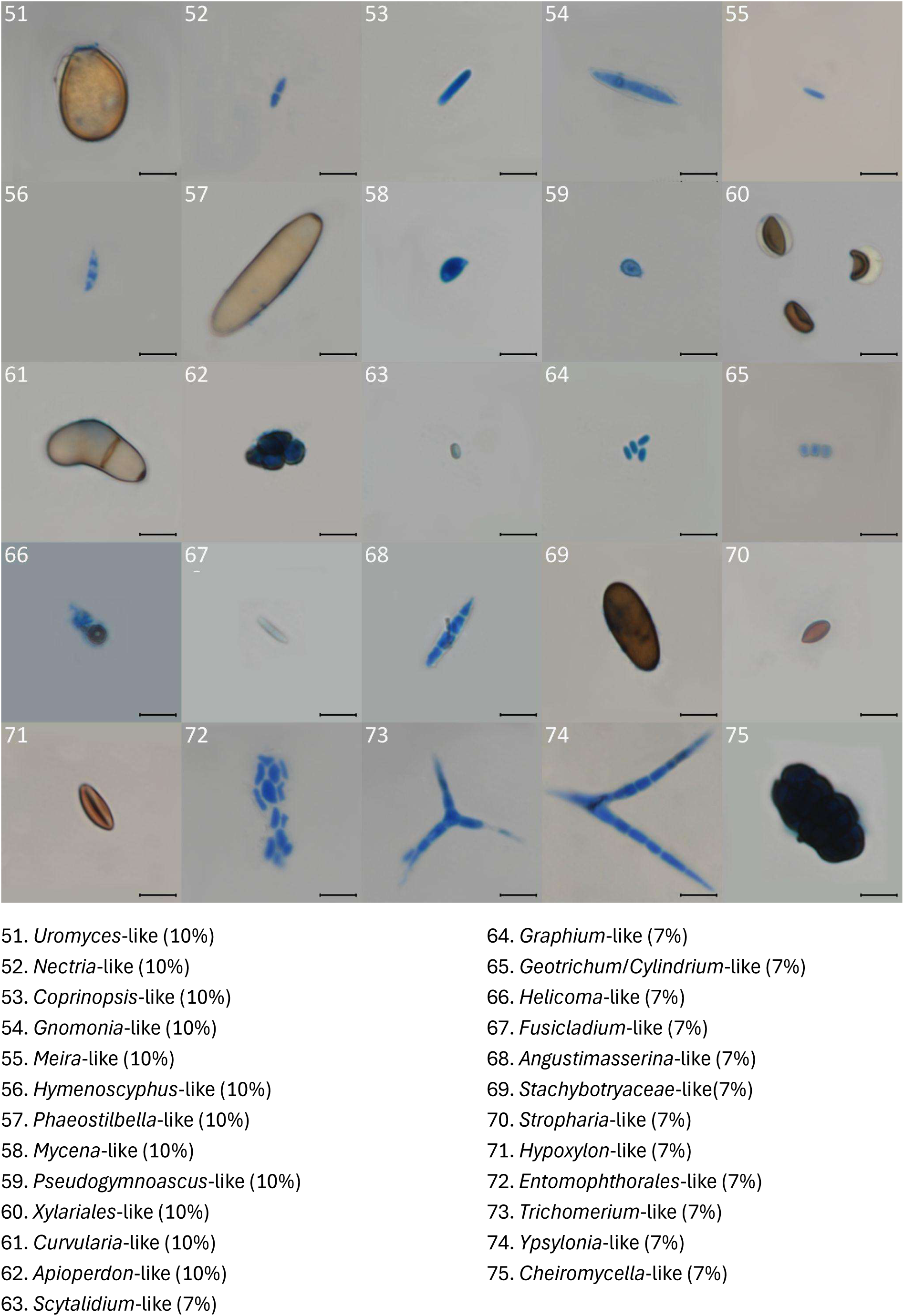
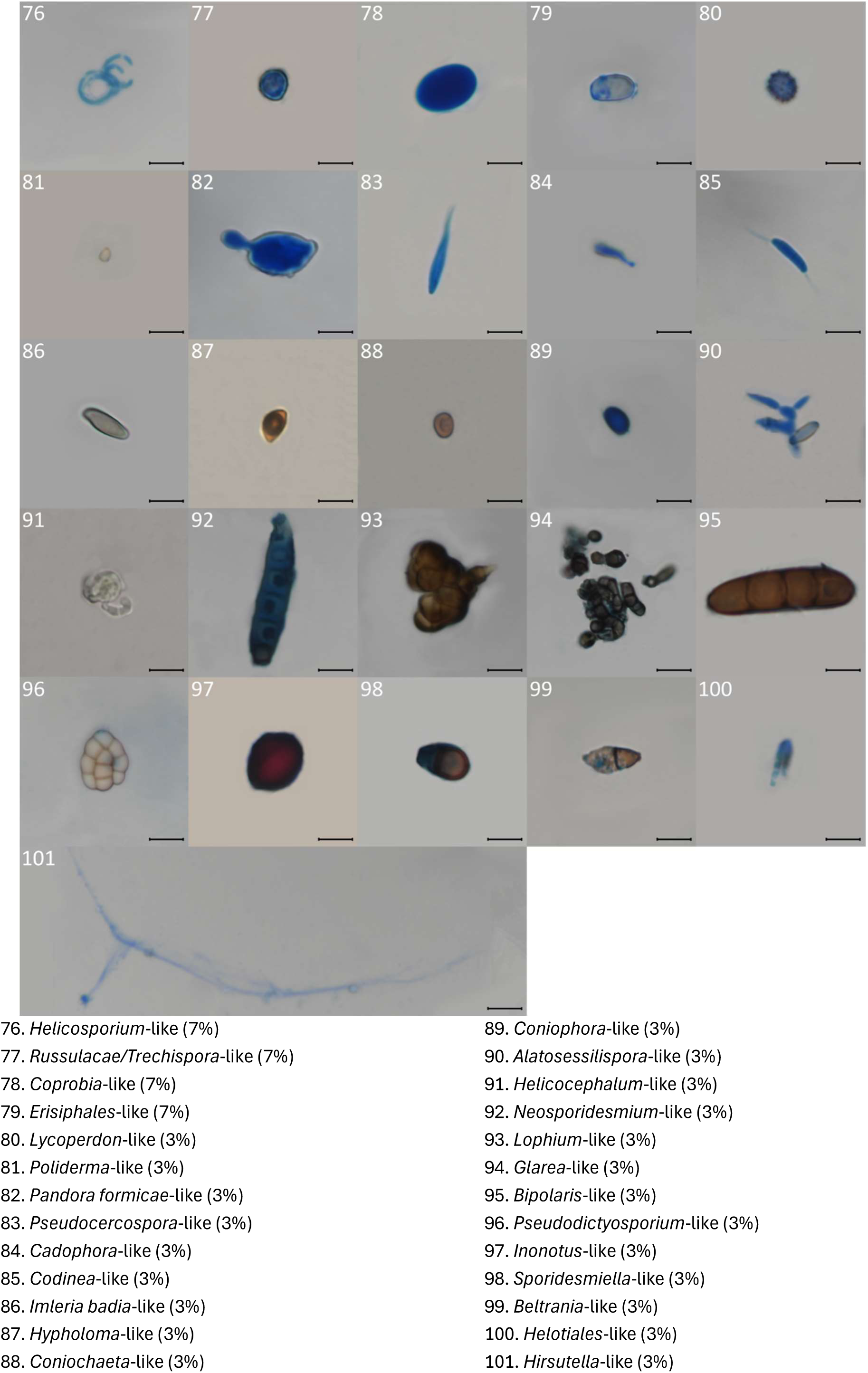

## A.4. Detailed information on marker genes and amplification reactions used in the study

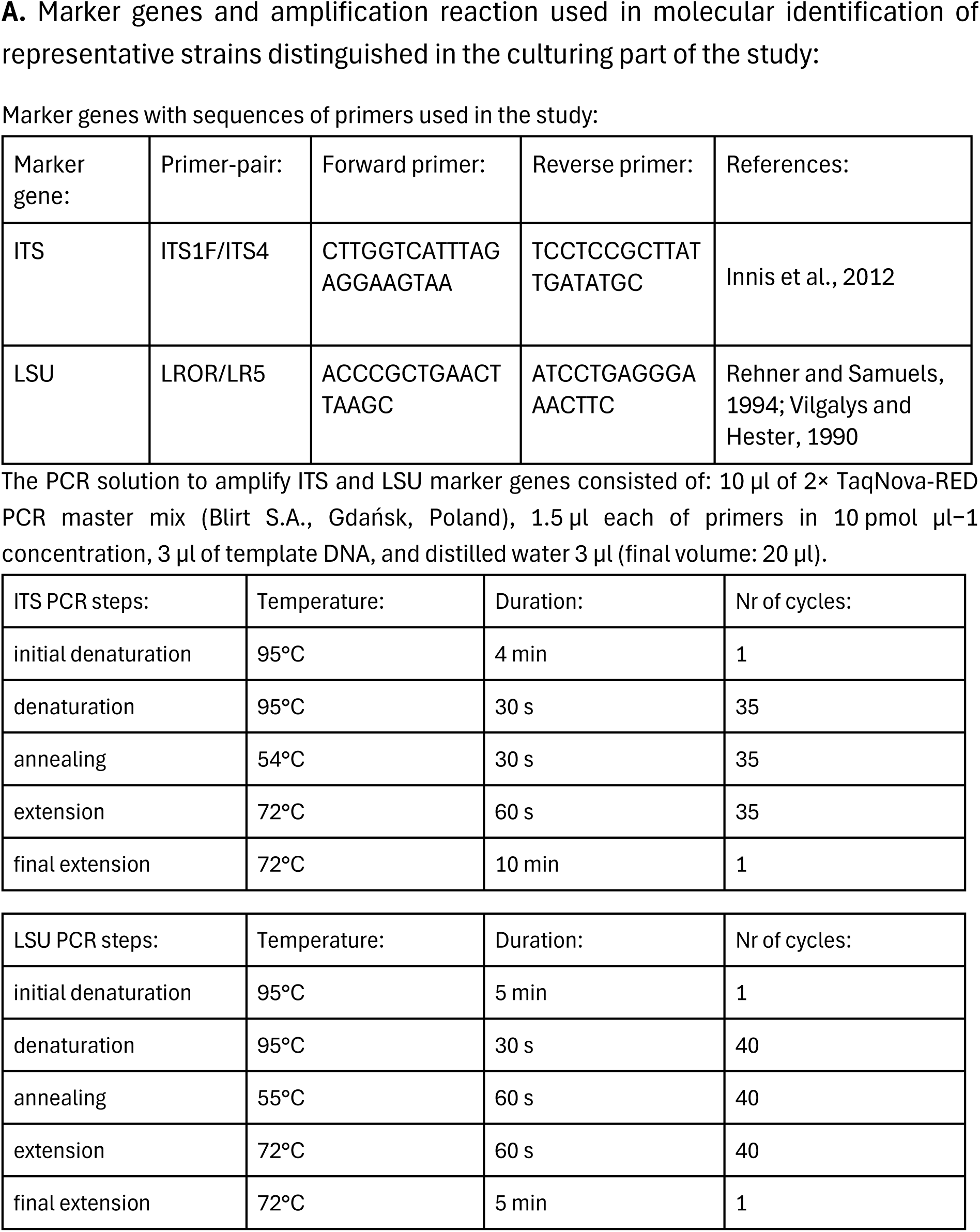
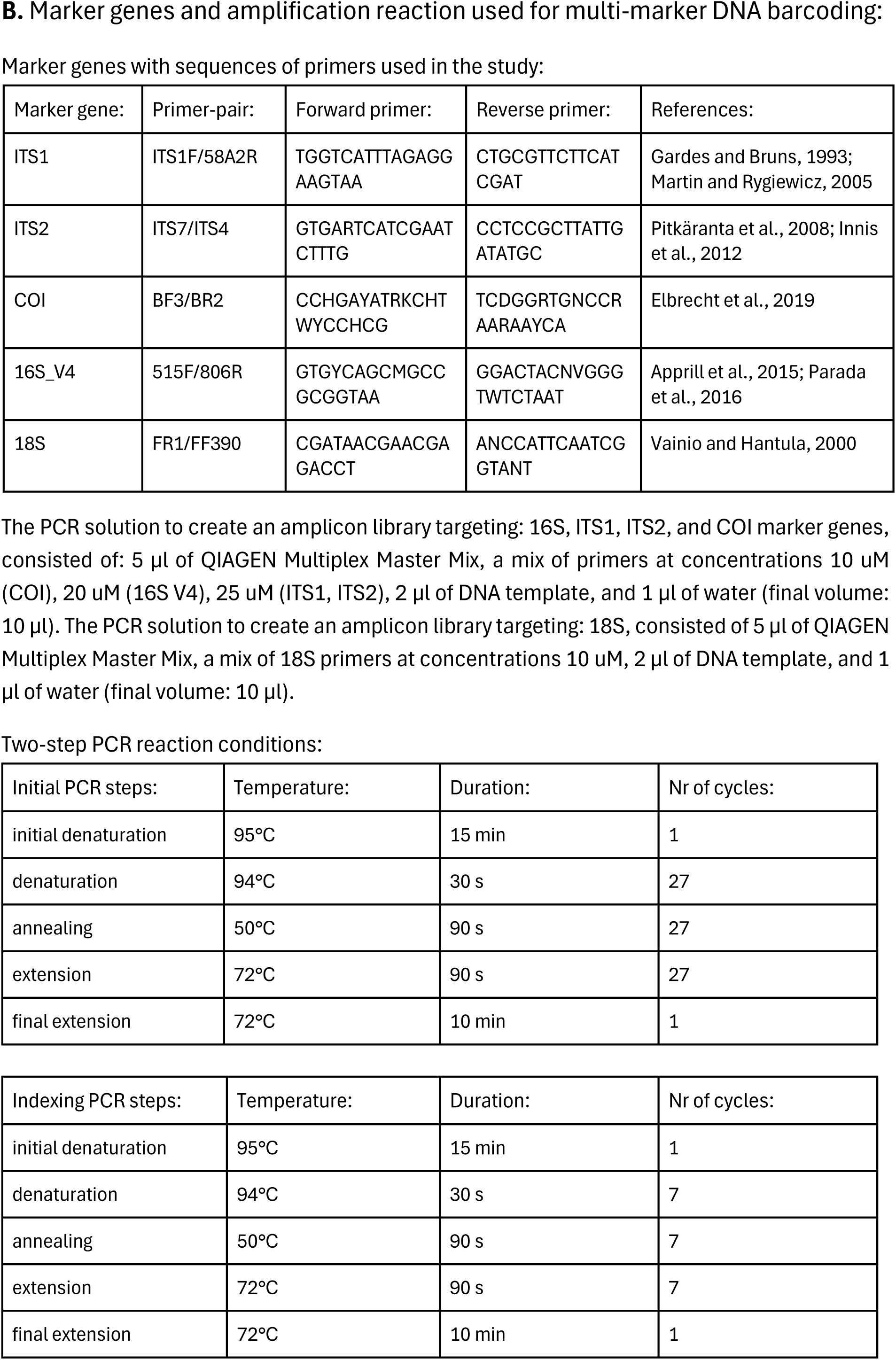

## A.5. Reference databases for taxonomic annotation of sequences obtained via metabarcoding

A.5. Reference databases for taxonomic annotation of sequences obtained via metabarcoding.
| Marker gene: | Database: | Version: | References: |
| --- | --- | --- | --- |
| ITS1 | UNITE | sh_general_release_dynamic_all_04.04.2024 | Abarenkov et al., 2024 |
| ITS2 | UNITE | sh_general_release_dynamic_all_04.04.2024 | Abarenkov et al., 2024 |
| 16S_V4 | SILVA | 138 SSU | Quast et al., 2012 |
| 18S | SILVA | 138 SSU | Quast et al., 2012 |

## A.6. Changes in taxonomy assignments for 18S reads

As taxonomy reference databases SILVA and UNITE are based on differing taxonomic backbones, taxonomy assignments for 18S reads were adjusted to be able to compare fungal diversity noted using the ITS marker gene. Following changes were made to the 18S dataset: for ASVs assigned as *Mortierellales*, the phylum assignment changed from *Mucoromycota* to *Mortierellomycota* (backed by Tedersoo et al., 2018); for ASVs assigned as *Cryptomycota*, the assignment was renamed to *Rozellomycota* (taxonomic synonyms); for ASVs assigned as *Cladosporiaceae*, order assignment changed from *Capnodiales* to *Cladosporiales*, and for ASVs assigned as *Mycosphaerellaceae*, order assignment changed from *Capnodiales* to *Mycosphaerellales* (backed by Abdollahzadeh et al., 2020); for ASVs assigned as *Buckleyzyma*, order assignment changed from Incertae Sedis to *Buckleyzymales* (backed by Zhao et al., 2017); and for ASVs assigned as *Hyphodontia*, order assignment changed from *Corticiales* to *Hymenochaetales* (backed by Wang et al., 2021).

## A.7. Genus-level taxonomic refinement of ASVs using BLAST

In this study, we accepted SINTAX-based taxonomic assignments for ASVs with a probability of at least 0.75 at the phylum level and at least 0.9 at all other taxonomic levels. However, to complement taxonomic assignment, ASVs that were not classified at the genus level and reached a maximum relative abundance of >1% in any library were compared with data available in NCBI GenBank (ncbi.nlm.nih.gov, accessed on 13 November 2024) using the BLASTN algorithm (Altschul et al., 1990). We assigned a genus-level name to an ASV when the top BLAST hits (restricted to type strains and reference strains deposited in recognized public culture collections) met the following three criteria: (1) sequence coverage >90%, (2) sequence identity >95% for ITS, >96% for 18S, and >99% for 16S, and (3) sequence identity to the closest other genus >1% for ITS and >0.3% for 18S and 16S. Otherwise, a higher taxonomic rank was used for ASV assignment.

## A.8. Detection and filtering of laboratory contaminants in metabarcoding datasets

To determine reads likely originating from laboratory contaminants in fungal metabarcoding datasets, we calculated the relative abundances of all ASVs across all samples and compared the maximum values of each ASV for experimental samples and in blanks. ASVs with an experimental-to-blank maximum value ratio of less than 10 were classified as likely laboratory contaminants and excluded from the analysis. However, as in the 16S rRNA dataset, we observed relatively high contamination rates (48% of reads in a library on average, see Figure below), we assigned contaminants at the OTUs level and excluded from further analysis all ASVs representing OTUs assigned as laboratory contaminants.

**Figure A.8.**
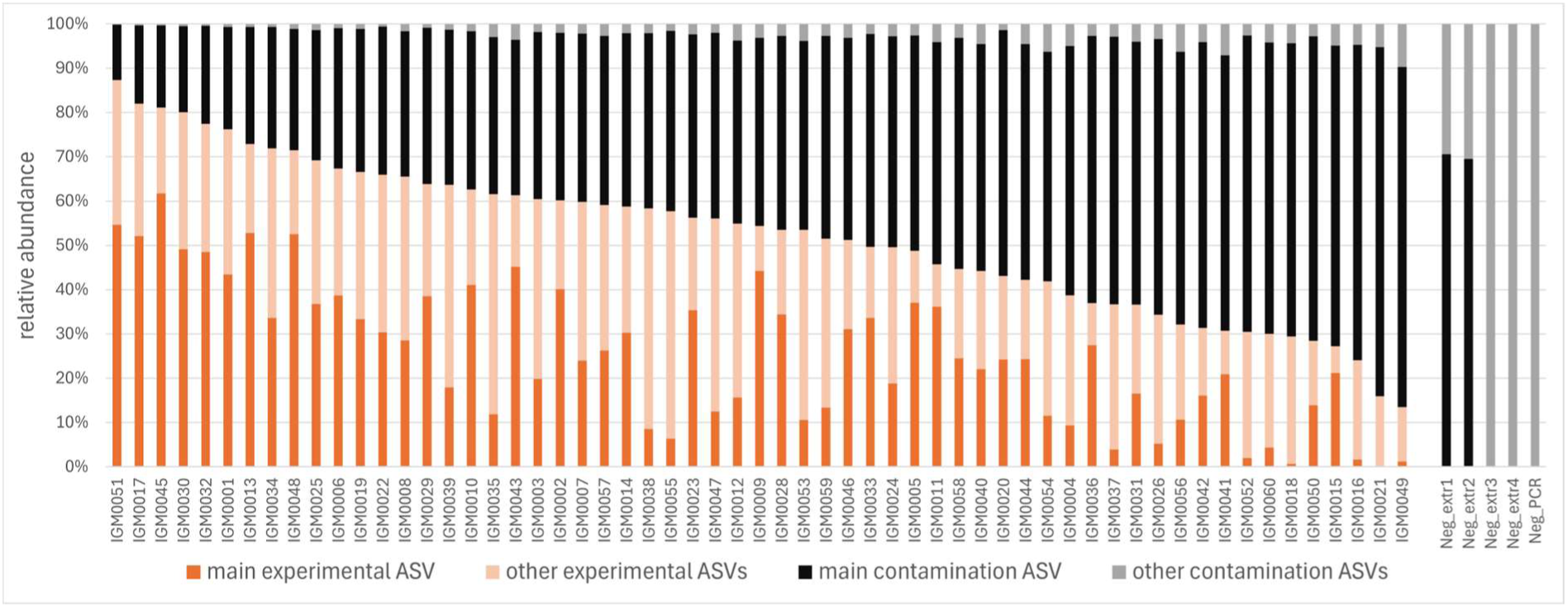
Contamination level of bacterial 16S rRNA libraries. Relative abundance of non-contaminant and contaminant ASVs in experimental and blank 16S rRNA libraries, with the most abundant non-contaminant ASV classified as a true ant associate (ASV_2 *Fructilactobacillus*) and the most abundant ASV classified as a contaminant (ASV_1 *Cellulosimicrobium*) highlighted. Experimental libraries are ordered by the descending relative abundance of non-contaminant ASVs.

## A.9. Correction of *Cladosporium* reads in 18S dataset

In all fungal marker gene datasets, genotypes assigned to *Cladosporium* were among the most abundant in experimental samples as well as blanks. However, in both ITS datasets, two abundant *Cladosporium* genotypes were present: one in blanks and experimental samples, and the other virtually exclusively in experimental samples (see below). We considered the first genotype as contamination and filtered it out. In the 18S rRNA datasets, offering much lower taxonomic resolution, only one *Cladosporium* genotype was present in both experimental samples and blanks. We concluded that this 18S genotype represents both reagent contaminants and true abundant symbionts. As a result, we decided to remove only a fraction of *Cladosporium* 18S reads, using ITS-based information on the proportion of total *Cladosporium* that the apparent reagent contaminant genotype comprises.

**Figure A.9.**
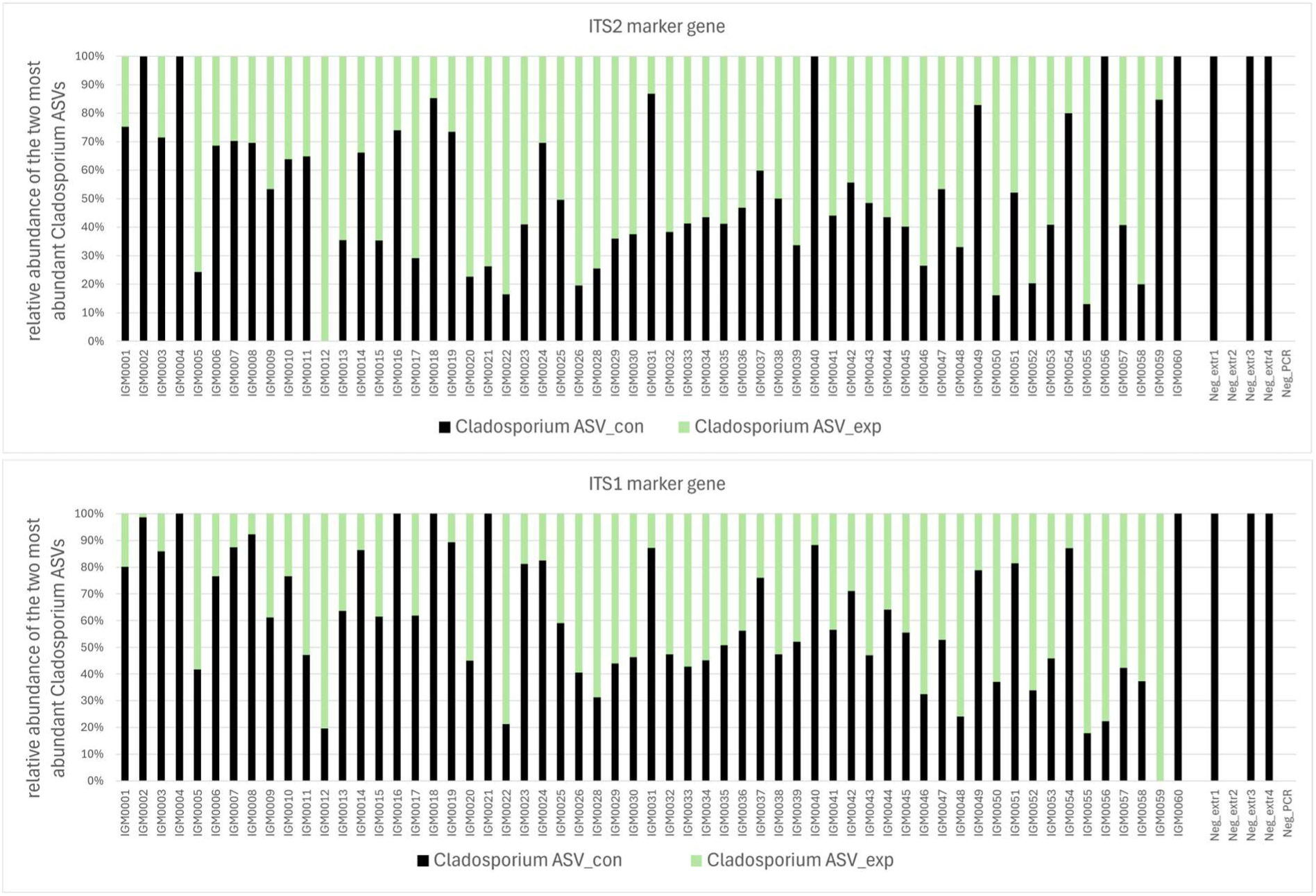
Relative abundances of the two most abundant *Cladosporium* ASVs in the ITS1 and the ITS2 datasets in experimental and blank libraries.

## A.10. Comparison of taxonomy assessment level in the ITS2, the ITS1, and the 18S metabarcoding databases

**Figure A.10.**
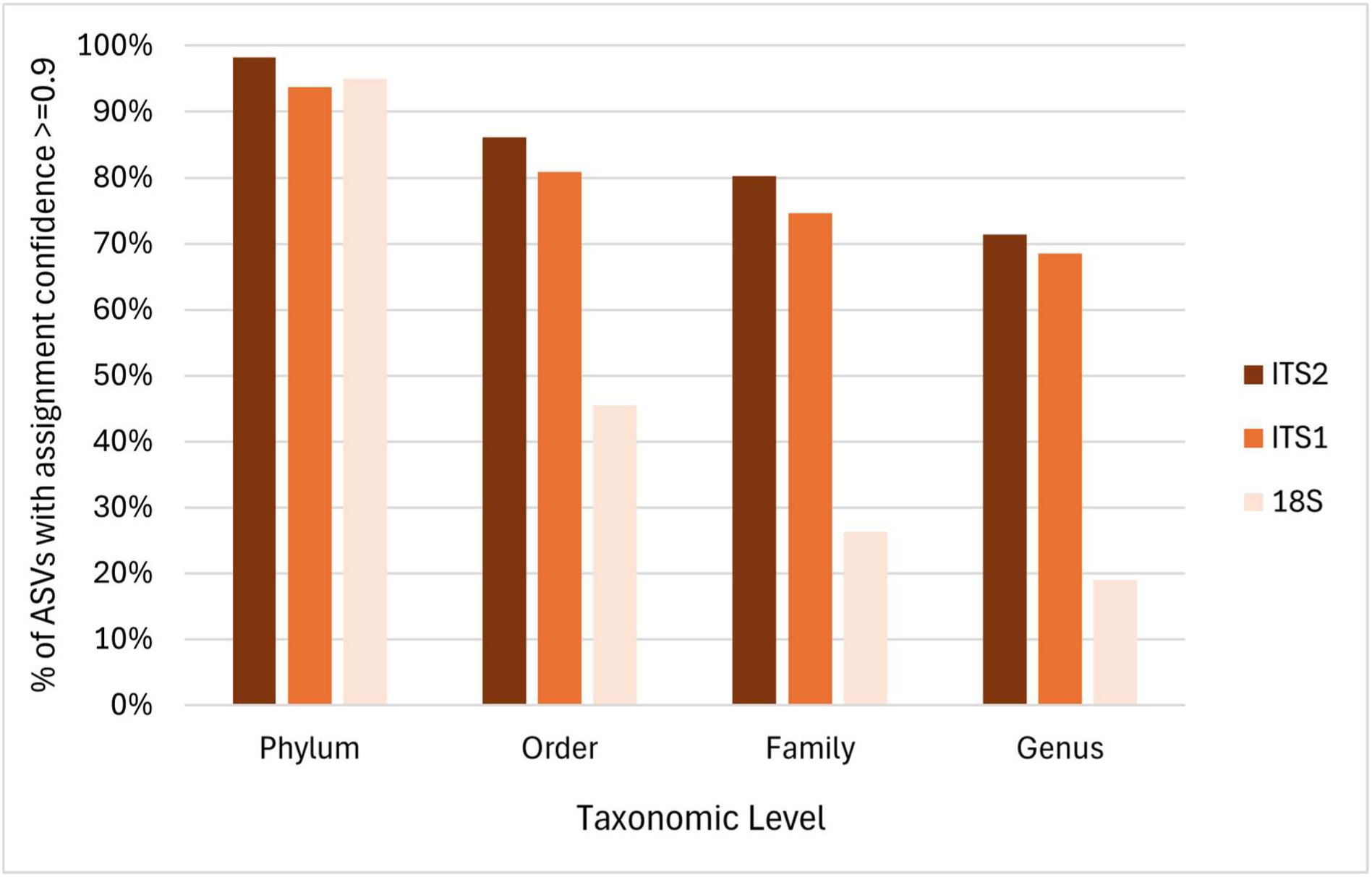
Taxonomy assessment level in the ITS2, ITS1, and 18S metabarcoding databases across different taxonomic levels (Phylum, Order, Family, Genus), estimated by the percentage of ASVs with high assignment confidence (p ≥ 0.9).

## A.11. Fungal ‘dark taxa’ diversity and distribution in the *F. polyctena* IBPs

**Figure A.11.**
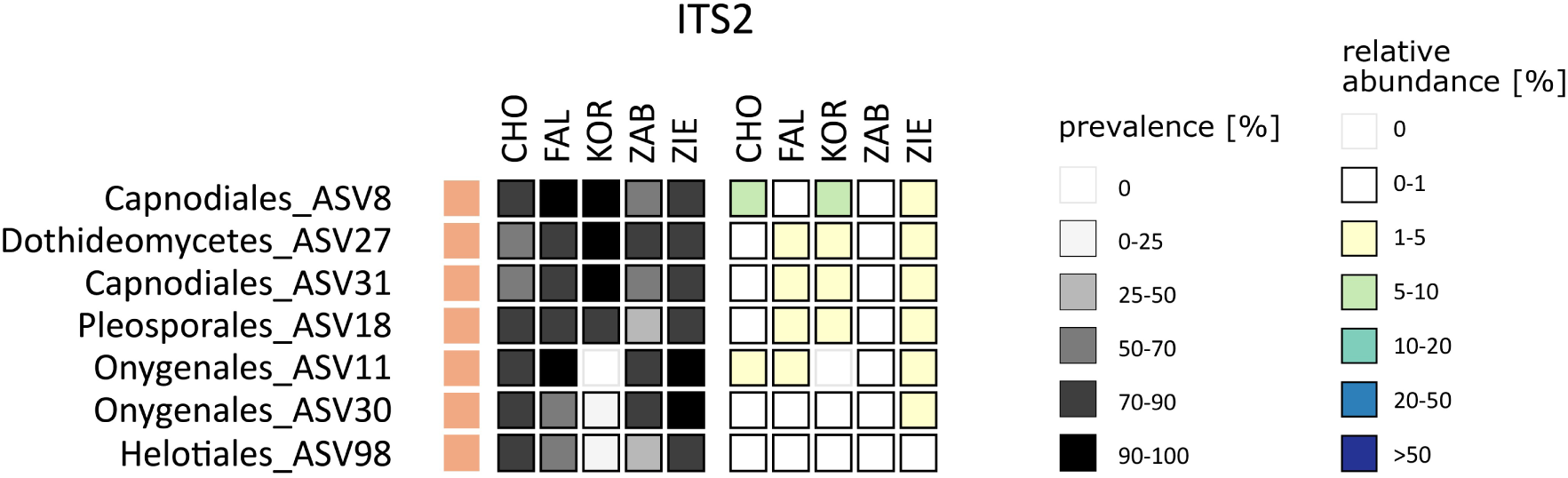
The distribution and average relative abundance across ant colonies of the most prevalent ITS2 ASVs, which lacked genus-level identification.

## A.12. Comparison of Sanger sequences obtained using the culturing approach with ITS2 and ITS1 metabarcoding datasets

**Figure A.12.**
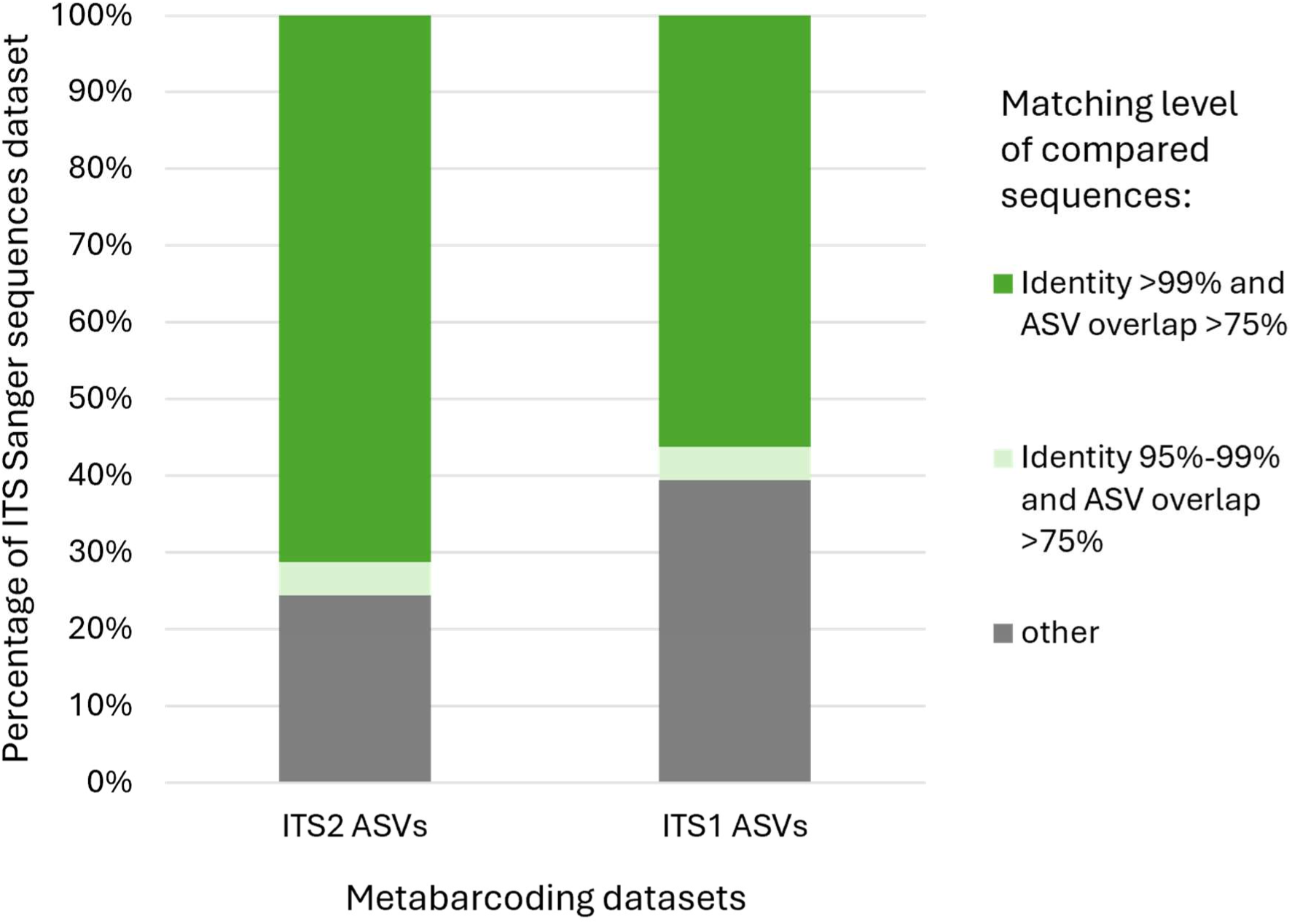
Completeness of ITS1 and ITS2 metabarcoding datasets relative to the ITS Sanger sequencing dataset.

**Table A.12.** Consistency of taxonomic assignment between Sanger ITS dataset and ITS1/ITS2 metabarcoding datasets. *Matching sequences requirements: identity >99%, ASV overlap >75%.

| Matching sequences* | Taxonomic assignment mismatch | Taxonomic identification uniform |  |  |  |
| --- | --- | --- | --- | --- | --- |
|  |  | type level | order level | family level | genus level |
| Sanger ITS vs. ITS2 | 0% | 100% | 98% | 98% | 85% |
| Sanger ITS vs. ITS1 | 0% | 100% | 92% | 89% | 82% |

